# Conformational variation in enzyme catalysis: A structural study on catalytic residues

**DOI:** 10.1101/2021.12.12.472283

**Authors:** Ioannis G. Riziotis, António J. M. Ribeiro, Neera Borkakoti, Janet M. Thornton

## Abstract

Conformational variation in catalytic residues can be captured as alternative snapshots in enzyme crystal structures. Addressing the question of whether active site flexibility is an intrinsic and essential property of enzymes for catalysis, we present a comprehensive study on the 3D variation of active sites of 925 enzyme families, using explicit catalytic residue annotations from the Mechanism and Catalytic Site Atlas and structural data from the Protein Data Bank. Through weighted pairwise superposition of the functional atoms of active sites, we captured structural variability at single-residue level and examined the geometrical changes as ligands bind or as mutations occur. We demonstrate that catalytic centres of enzymes can be inherently rigid or flexible to various degrees according to the function they perform, and structural variability most often involves a subset of the catalytic residues, usually those not directly involved in the formation or cleavage of bonds. Moreover, data suggest that 2/3 of active sites are flexible, and in half of those, flexibility is only observed in the side chain. The goal of this work is to characterise our current knowledge of the extent of flexibility at the heart of catalysis and ultimately place our findings in the context of the evolution of catalysis as enzymes evolve new functions and bind different substrates.

## Introduction

The “structure determines function” concept is fundamental for proteins and especially in enzymes, whose catalytic potential is largely dependent on its active site 3D shape. Active sites of similar enzymes are essentially conserved in sequence[1,2] and structure, however geometrical variation is commonly observable[3]. Evolutionary unrelated enzymes can converge towards similar functions with their active sites adopting similar geometries[4,5], while the outer shell of the protein is freer to vary.

Variation in active site conformations can be described by two terms: plasticity and flexibility[6]. Plasticity refers to differences between related enzymes during the evolution of the active site geometry whilst retaining the function, while flexibility refers to the movement of the functional groups within the active site in an individual enzyme. In a study by Todd et al. (2002)[6], various examples of active site plasticity are reported, and these cases are classified according to the observed type of structural change. Phenomena like circular permutations[7], residue functional substitution, convergent evolution and divergence followed by convergence[4] are some of the mediators of variation within enzyme families and superfamilies.

Flexibility can be captured as alternative conformations in different crystal structures of the same protein. Such conformational shifts are usually driven by ligand binding, with enzymes often adopting an “open” form before the substrate enters the active site (-apo or resting state) and a “closed” form being stabilised upon substrate binding. This model of behaviour was captured by the “induced fit” theory, described by Koshland[8], which “replaced” the “lock and key” model of ligand binding, where the active site of an enzyme would be structurally compatible with a single type of substrate and would not exhibit significant flexibility. It has been shown that ligand binding sites can be flexible to different degrees, and the magnitude of variation is related to the variation of the ligands able to bind in the site[9]. Active site flexibility is essential for catalysis to initiate and proceed[10], particularly in the context of enzyme-ligand interactions. Previous studies have presented several cases of flexible active sites[11,12] and have distinguished different types of motion in the catalytic residues. Similarly, a systematic study by Weng et al.[13] followed an algorithmic approach for the identification of flexible active sites in oxidoreductases and the quantification of their flexibility.

Many structural surveys on enzyme active sites have used a common data resource to collect manually curated catalytic residue annotations - the Mechanism and Catalytic Site Atlas (M-CSA)[14] (or its precursor, the Catalytic Site Atlas – CSA[15,16]). With the vastly growing number of available protein structures in the PDB, we are now able to perform a highly comprehensive and systematic analysis on conformational variation in enzyme active sites, involving more than 60,000 structures and annotations for 925 diverse enzymes families covering a large portion of the functional space, giving the broadest to-date structural survey on active sites. A dynamic superposition method is implemented to align and compare active sites from homologous enzyme structures, providing the ability to quantify conformational differences at single-residue level. We attempt to show relationships between sequence and structure within evolutionary related enzymes, and by introducing explicit annotations for all ligands in proximity to active sites, we are able to characterise their impact on the geometry of the active site. This work in its broader context aims to provide a detailed structural description and characterisation for all enzyme families curated in M-CSA, something that can be utilised in enzyme repurposing and design.

## Results

### Sequence and structural similarity relationship in enzymes

Enzymes from homologous families within M-CSA were compared by their sequence identity (expressed as a percentage) as well in their structural similarity, both on the whole protein structure and on the isolated active site. For structural alignment of active sites, this was performed either over their C_α_ atoms or their functional atom triads, with the relevant definitions being presented in Fig S1. In Fig. 1, we present an ensemble of scatter plots where each point represents a comparison between the reference enzyme of each M-CSA family and a member of its homologues (See Methods section for dataset details). Each panel represents potential relationships between different metrics; the correlation was assessed by linear regression and Pearson correlation coefficient calculation.

**Fig. 1:**
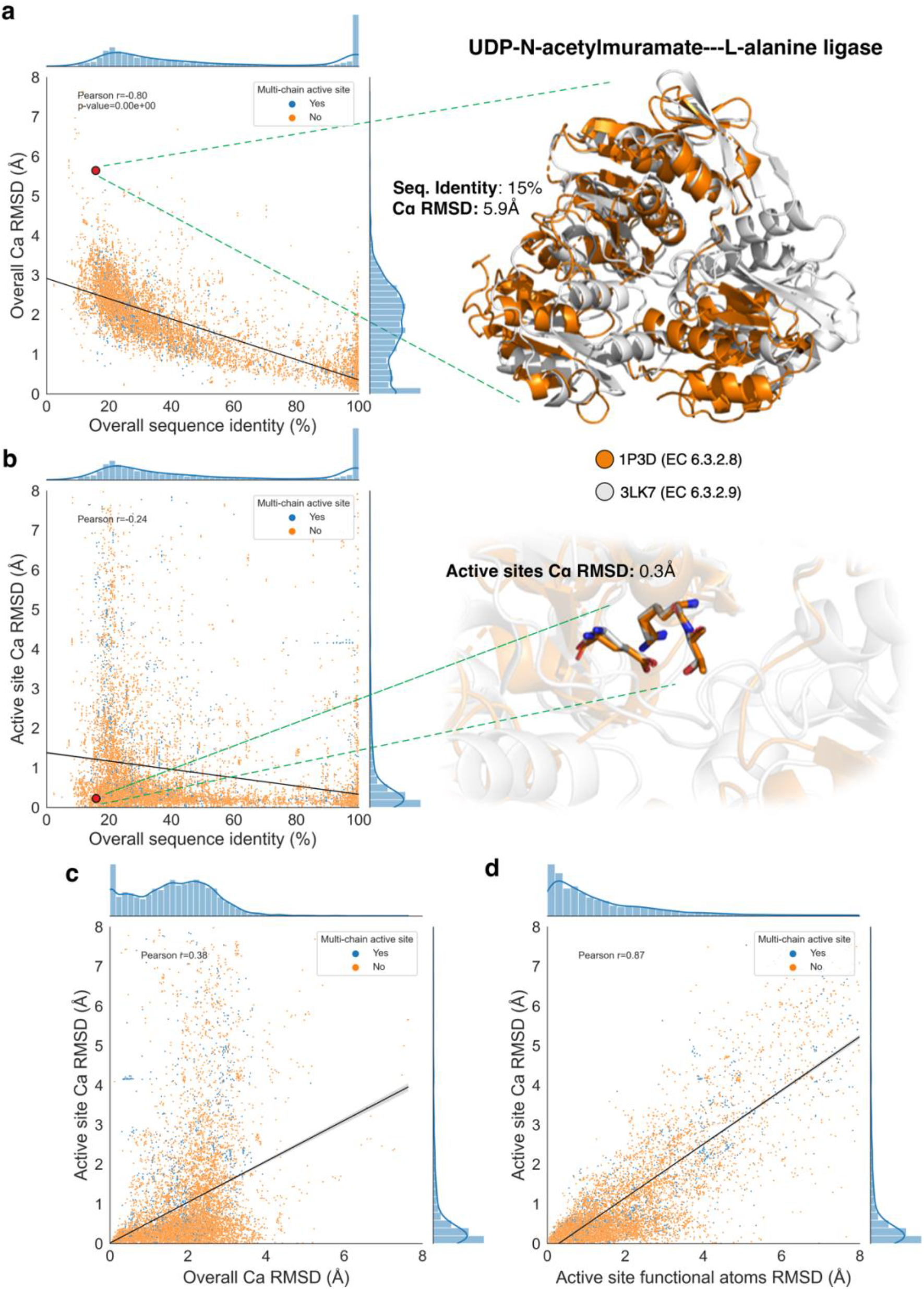
Pairwise comparison of the reference enzyme in each M-CSA family with all its homologues, in sequence and structure (30,859 data points in each plot). a and b: Left part of each panel shows the percentage of overall sequence similarity vs. RMSD over all C_α_ atoms (a) and the percentage of overall sequence similarity vs. RMSD over active site C_α_ atoms (b). Right part illustrates an example comparison of two homologous enzymes in superposition over all C_α_ atoms (a) and over the C_α_ atoms of their active sites (b). The corresponding data point on the plots (left part) are indicated by red circles. c: RMSD over all C_α_ atoms vs. RMSD over active site C_α_ atoms. d: RMSD over active site functional atoms vs. RMSD over active site C_α_ atoms (refer to text for functional atoms definitions). In all plots, data points corresponding to enzymes whose active site is composed of residues belonging to multiple chains, are indicated by a different colour and the distributions of data points on each axis is shown as inline histograms. 3D structure models were prepared in PyMol[18].

The first type of relationship explored was between the overall sequence similarity and overall structural similarity, expressed as the RMSD over the C_α_ atoms of the whole protein. As shown in Fig. 1a, an increase in sequence similarity is accompanied by an increase in overall structural similarity (Pearson *r=-0.81, p*-value≃0), a trend that has been described extensively for multiple types of proteins[5],[17]. However, a marked shift of this sequence-structure trend can be seen when focusing on the structural similarity of the active site instead of the overall structure. Fig. 1b demonstrates that if we only consider the alpha carbons of the active site residues, this relationship between sequence and structural similarity is broken. The distribution of data points (shown as inline histograms) exhibits high density in low active site RMSD values (i.e. <2Å) compared to the overall RMSD values that are distributed more evenly up to ~4Å. These distributions indicate the high structural conservation of the active site, almost regardless of the sequence similarity. Several examples exist that are consistent with this hypothesis, with one illustrated in the right part of Fig. 1a and Fig. 1b: Here, two distant homologues of the L-alanine ligase family (M-CSA 876), dissimilar both in sequence (15% identity) and in overall structure (C_α_ RMSD=5.9Å), but similar in function (EC number differing only on the fourth level), share an almost identical conformation in their active sites (0.3Å), even though one of three catalytic residues is not conserved. In this case, the catalytic centre remained conserved in its conformation during evolution, while the outer shell of the protein was less subject to evolutionary pressure, therefore, was freer to vary. Another interesting observation in the plot of Fig. 1b is that high structural dissimilarity values can even be observed at almost sequence-identical enzyme pairs, suggesting that other factors, such as ligand binding and environmental conditions, can influence the conformation of active sites significantly and in various degrees. This prompted up to investigate the role of ligands in active site structural transitions, which will be discussed further below.

The conserved nature of active sites on the structural level, as well as their flexibility, can also be observed when looking at the relationship between the active site and the overall structural dissimilarity (Fig. 1c). It can easily be seen that even in high RMSD values (up to 4Å) on the whole structure, active site RMSD remains low (<2.5Å). Nevertheless, the conformation of active sites can considerably vary, even in structures that share high structural similarity overall.

Finally, we compare the backbone of the catalytic residues (C_α_ atoms specifically) and the atoms participating in the reaction, which most often belong to the side chain. Plotting these metrics –the active site functional atoms vs. C_α_ RMSD (Fig. 1d)– we see that although there is undoubtedly a linear relationship between them (as expected), that backbone atoms are more constrained than the functional atoms during catalysis, and most of the flexibility in catalytic residues derives from the side chain. We can define as “rigid” and “flexible” active site pairs, the ones under and over an RMSD threshold of 0.5Å (limit of experimental error). Using this definition, 32% of active sites in this dataset are rigid both in main and side chain, 35.6% are flexible both in main and side chain, 32.1% only exhibit side chain flexibility and a small 0.3% fraction is rigid on side chain, but main chain atoms are flexible. Therefore, the general observation out of this is that ~2/3 of active sites exhibit some degree of flexibility, and in half of those, this flexibility is accounted only to side chain motion.

It should be commented here that the *p*-values of the Pearson correlation coefficients in all four analyses presented are close to zero, something that can be accounted to the high number of observations in our dataset. This provides confidence that the trends presented in these results are reliable, especially after the removal of artefacts, as described in the Methods section. Furthermore, we distinguish pairs of single-chain and multi-chain active sites which occupy ~89% and ~11% of our data respectively (data points coloured differently in the plots). Here, there is no apparent pattern in the distribution of data corresponding to multi-chain active sites, which indicating no significant difference in the structural behaviour between single and multi-chain enzymes. This is further supported by the individual Pearson correlation coefficients for these two types, which are similar in all cases (values not shown in the figure).

### Effect of ligands in active site structures

One of the main goals of this work was to assess the impact of ligands binding to the active site, with respect to changes in the conformation of the catalytic residues. For this purpose, we performed an all-vs.-all structural comparison of the active sites within each homologous family in M-CSA, using the weighted superposition algorithm over functional atom triads, as described in the Methods section.

Data were split into two groups: (i) active sites that derive from the same, or (ii) different protein sequence (by checking identity in their UniProt mapping). Each group was further split down to subgroups of comparisons in which active sites have the same or different catalytic residues (two structures can have the same UniProt mapping but catalytic residues might be mutated). The final grouping was based on the presence of native-like ligands in the active site. We distinguish three categories: (i) both homologous active sites are ligand-free, (ii) both are bound with at least one native-like ligand (≥60% PARITY score with a native enzyme reaction component[19]), and (iii) one site is ligand-free and one is ligand-bound. For each M-CSA family, the average RMSD for each group was calculated and the distributions of these average RMSD values are presented in Fig. 2. Fig. 2a and 2b show that active sites deriving from the same protein exhibit lower RMSD values (tend to be more similar) than those deriving from different proteins. However, in both these subgroups, when distinguishing between site pairs with and without catalytic residue mutations, we see the highest RMSD shift, with mutations being associated with significantly higher RMSD values. More subtle yet significant conformational changes occur due to perturbations by ligand binding: here, the highest structural similarity is observed when both active sites bind at least one native-like ligand (“Bound vs. Bound” groups). In contrast, ligand-free active sites tend to be more variable (“Free vs. Free” groups), and the highest variability is observed when comparing between ligand-free and ligand-bound active sites (“Bound vs. Free” groups). It should be noted that the most significant conformational differences among these three subgroups can be found in conserved active sites from the same protein (Kruskal-Wallis test[20] *p*-value=2.12×10^−6^), while the least significant differences are in mutated active sites from different proteins (Kruskal-Wallis test *p*-value=0.16). These shifts are also demonstrated by linear regression of mean RMSD (shown as a red line over the boxplots), with the slope of the regression lines reflecting the degree of significance. The main suggestion here is that active sites are flexible in their apo-form and upon binding of a ligand, structural stabilisation occurs. Moreover, the median value in the distribution of RMSDs in “Bound vs. Free” active site pairs is slightly lower than in “Free vs. Free” ones, indicating that ligand-free active sites often adopt the catalytically active conformation. This further suggests that during its pre-binding structural transitions, an active site can pass through a “primed” conformation, favouring substrate binding. This claim is supported further below in our case studies.

**Fig. 2:**
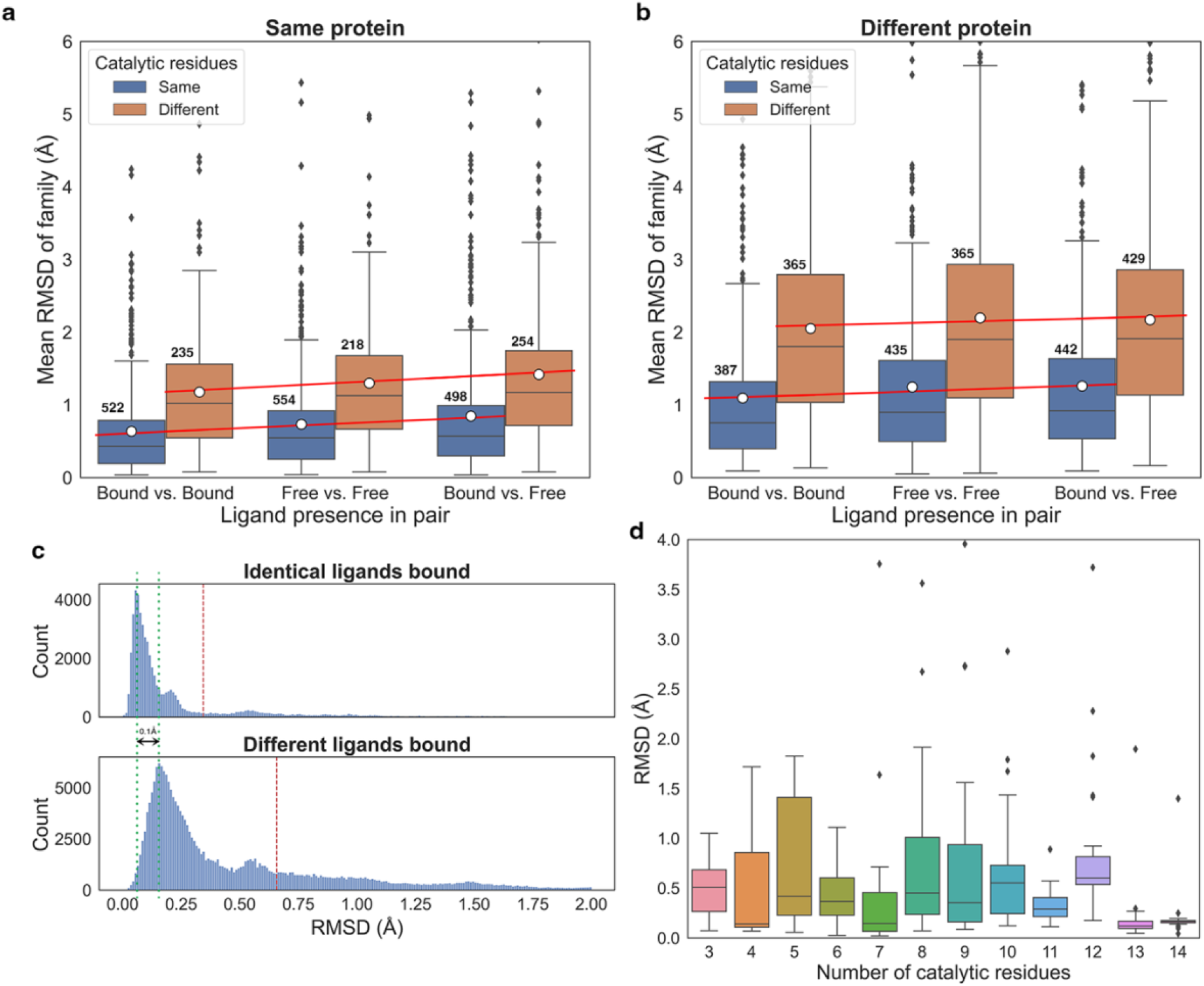
Conformational variation between active site pairs of homologous enzymes, expressed as the Root Mean Square Deviation of their functional groups’ atomic positions. a and b: Distributions of the average pairwise active sites RMSD of homologous enzyme families, grouped by the presence of ligands in each active site comparison pair. Panels a and b refer to pairs of enzymes that have the same or different UniProt mapping respectively. Active site pair groups that have the same or different catalytic residues (e.g. by mutations in the PDB structure) are colored differently (see legend). The mean and median RMSD values are marked by a horizontal line and a white circle respectively. Simple linear regression to show differences on the mean of each group is drawn as a red line. Size of each group is labelled above each boxplot c: Distributions of functional atoms RMSD when the same (upper panel) or different ligands bind to the active site (lower panel). The dataset used in this plot includes only enzymes that have identical UniProt mapping and the same catalytic residues. Red dashed lines indicate the mean RMSD value, while green dotted lines indicate the RMSD values of the major peaks, with their difference being labeled. d: Distributions of functional atoms RMSD of ligand-free active site pairs from the same protein and with the same catalytic residues, grouped by the number of catalytic residues.

Although this overview is informative and can give some insight on the influence of ligands in the active site, some caveats exist. The first regards the criterion to define an active site as “ligand-bound”: our definition requires at least one non-artefact, native-like ligand to be present close to the catalytic residues without specifying its type (substrate, co-factor, or ion) and the overall number of ligands that bind. This could alter the average active site RMSD, especially in families with a few homologues, as different ligands can cause different conformational changes. This can particularly be seen in Fig. 2c, where we demonstrate the impact of binding the same or different ligands in identical active sites from identical proteins. We see that there is significant increase in the mean RMSD when different ligands are bound, whereas binding of the same ligands usually means highly similar conformation (we consider RMSD values below 0.5Å to fall within experimental error). However, significant conformational differences can also be seen even when the same ligands bind, something that supports the notion of flexibility during catalysis – i.e., an active site might be captured in different conformational states during crystallisation, even when the same ligand is introduced[21]. Another caveat is the presence of low-populated enzyme families in the dataset; these could influence the mean RMSD value as there are less observations to be averaged out. Furthermore, another concern in this analysis was the impact of the number of catalytic residues to the active sites RMSD. Fig. 2d shows distributions of functional atoms RMSD, when the comparison pairs are grouped by the number of catalytic residues. Here, there is no evident correlation between the number of catalytic residues and RMSD, therefore, we feel that this would not introduce significant bias. Last, a comment should be made on outlying observations in Fig. 2a. These outliers, as discussed above, could be accounted to small enzyme families, or to highly heterogenous families, or even to superposition artefacts. Although the active site superposition algorithm is robust and optimised, extreme cases do exist, where RMSD is unusually high. For instance, in some cases, 3-residue active sites might appear flipped in the superposition, because their geometry is such that during fitting iterations, one residue might be excluded, and the other two can be superimposed ambiguously. Although these cases are very few and additional filters were implemented to exclude them, some remains do still exist and their number is boosted due to the all-vs.-all nature of this analysis.

Finally, it should be noted at this point that this is just a coarse overview, nevertheless it provides a general insight on active site variation and flexibility, as well as in the role of ligands to active site conformation. Enzymes are highly heterogenous and not all families are the same in terms of sequence and structural variability. Moreover, as will be discussed further below in specific case studies, the global RMSD over a subset of atoms of the catalytic residues is only an approximate way to screen for difference in active site conformation. It does not provide any spatial information, that is, which and how many residues are flexible, and can be averaged down in cases where, for instance, a single residue is flexible in a large active site. This, in addition with the fact that different enzyme families exhibit different degrees of catalytic residue flexibility, makes it essential to view each family separately, and aggregation should be performed with care.

### Residue flexibility and functional roles

Given the conformational variation of an active site, does this variation reflect the role of the residue in catalysis? After extracting individual residues and their functional atoms RMSD values from the all-vs-all active site superpositions dataset we performed an enrichment analysis on residue type, functional role, and broader role category using annotations from M-CSA[2].

Fig. 3a and Fig. 3b present collections of single-residue RMSD distributions for each residue type and role respectively. Most residues, regardless of their type and role, remain relatively immobile with their RMSD lying within limits of experimental error (~0.5Å). This means that either the propensity to be flexible or rigid is unrelated to catalytic residue function, or any existing preferences are subtle and difficult to be distinguished in this representation. To address this, we compared the enrichment frequencies in two subgroups, comprising either “rigid” or “flexible” residues with the former being defined as the ones with RMSD<0.5Å and the latter with RMSD≥0.5Å. These results are presented in Fig. 3c-e. The “flexible”-over-”rigid” odds-ratio of the occupancies corresponds to the likelihood of a residue type/role/category to be flexible over rigid. For residue type, it is evident that positively charged residues (Lys, Arg) tend to be more flexible compared to negatively charged ones (Asp, Glu) (they are also longer) while the zwitterionic His exhibits the highest rigidity. This last observation is consistent with the fact that His is the primary residue involved in metal ligand binding[22]. Residues of the “reactant” category (proton donor, acceptor, and shuttle) appear as considerably more rigid (Fig. 3c) compared to “spectator” and “interaction” residues. This is justified in physical-chemical terms, as unnecessary flexibility at the reaction centre would lead to increase of entropy, lowering reactivity and catalytic activity. Conversely, “spectator” residues (electrostatic stabiliser, activator, steric role) are more flexible. The two positively charged hydrophilic residues, Lys and Arg, usually lie in the outer layers of the active site, participating in interactions with bulk solvent and in charge stabilisation.

**Fig. 3:**
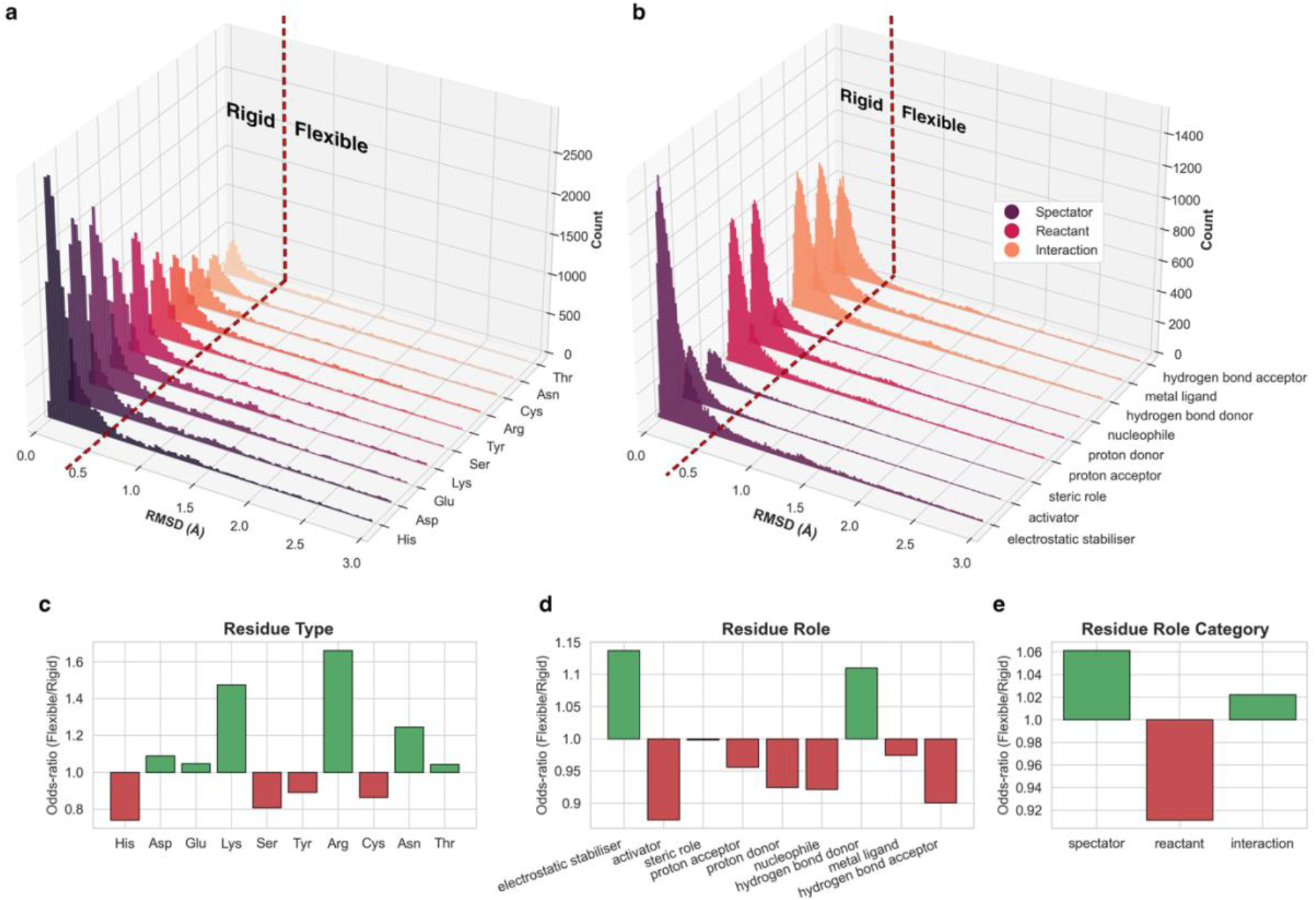
Enrichment analysis of conformational variation according to residue type and functional role in pairwise active site superpositions dataset. a and b: Single residue RMSD distributions in various residue types (a) and functional roles (b). Red dashed lines indicate the 0.5Å RMSD cut-off to define residues as “rigid” or “flexible”. c-e: Ratios of enrichment frequencies of each residue type (c), functional role (d) and role category (e) in two subsets of residue RMSD observations, one containing only residues defined as “rigid” and the other only the “flexible” ones. The odds-ratio of the frequencies represent the propensity to be flexible or rigid. Ratio >1 indicates that this specific type/role/category tends to be more variable in the 3D space, thus more flexible, while <1 indicates higher potential to be rigid. The dataset used here is sampled (50 superpositions of conserved and identical active sites from the same protein per M-CSA homologous family). After enrichment and grouping, only groups occupying at least 2% of the sampled dataset are reported.

Although hydrogen bond donors and acceptors constitute a relatively small fraction of the dataset (12% and 7.6% respectively, out of 413,802 data points), acceptors tend to be more flexible, while donors are rigid. It is well known that the geometry of a donor is more restricted that that of an acceptor, and this is reflected in our observations[23].

Last, we analysed potential relationships between single residue RMSD and physical-chemical parameters such as B-factor, solvent accessibility, hydrogen bonding and residue-ligand contacts, with these results presented as correlation scatter plots in Fig. S2. We observed that RMSD values increase as the B-factor and solvent accessibility of the reside increase; in contrast the RMSD values decrease as the number of hydrogen bonds or ligand contacts increase. These results are to be expected but correlations are weak (Pearson r = ~0.2), probably due to the huge variation between individual residues and the different magnitudes of active site flexibility among enzyme families.

### Inherent rigidity and flexibility

Active sites can exhibit flexibility to different degrees with different fractions of their catalytic residues varying in 3D location. For each family, for which there are sufficient data (at least 50 homologous active site structures (307 in total), we generated a series of functional atoms RMSD distributions between all members of a given family. From these results we identified four different structural variability patterns in active sites (See Figs. S3 & S4), exemplified by several cases of enzyme families (3D superposition models in Fig. 4a and Fig. 4b). Visual inspection of the RMSD distributions of Fig. S3 provided an overview of the structural behaviour of the active site in each homologous enzyme family (excluding all structures with catalytic residue mutations in this case).

**Fig. 4:**
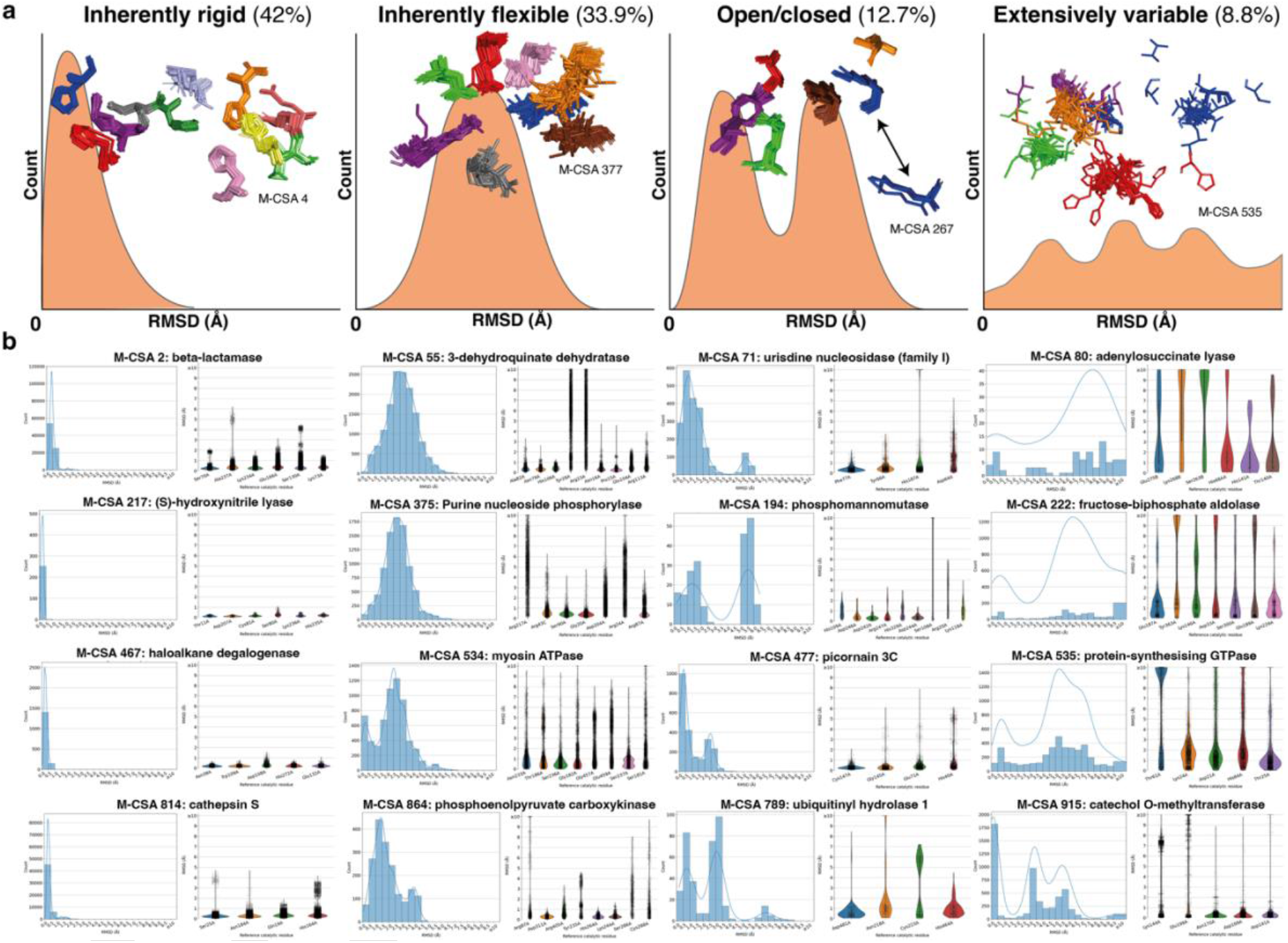
Different structural paradigms of active site structural behaviour (inherently rigid, inherently flexible, open/closed and extensively variable) (presented here in orange schematic histograms). For each paradigm, conserved active sites from a representative M-CSA enzyme family are shown in superposition, with the M-CSA identification number of each example entry being annotated. Two distinct active site conformations in the Open/closed type are indicated by a bidirectional arrow. 3D models were prepared in PyMol[18]. b: Examples of M-CSA families adhering to each paradigm. Plots on the left are histograms of all-vs-all functional atoms RMSD over the whole active site, and on the right, the RMSD distributions are plotted on a per-residue basis. Only conserved active sites were used to generate these distributions.

We are cognizant of the fact that our data set is limited – for example with different numbers of different types of ligands for different families. Here we have analysed the distributions we have been able to generate and extracted some “paradigms” as an aid to understand and describe these distributions. It should be noted that residues in loops often have poor electron density and therefore the observed structural variation might be largely ascribed to lack of experimental data. Therefore, given that our approach uses snapshots of conformational states in crystal structures to characterize 3D variation, the term “flexibility” used herein has a specific limited meaning.

With these caveats in mind, we distinguish four major active site structural paradigms in the context of flexibility, presented schematically in Fig. 4 and accompanied by four example M-CSA families (Fig. 4b): (i) **Inherently rigid**, where catalytic residue flexibility lies within experimental error. These “rigid” catalytic sites follow a left-skewed RMSD distribution, with a mean RMSD value usually below 1Å. A representative example (leftmost panel of Fig. 4a) is the family of nitrite reductases (M-CSA 4, EC: 1.7.2.1, reference PDB ID: **1NIA**) in which catalytic residues cluster perfectly without significant variability. Families of this type are abundant and constitute about 42% of the 307 entries. (ii) **Inherently flexible**, where a large fraction of catalytic residues tends to be flexible, each one to varying degrees. RMSD distribution here resembles a Gaussian distribution, usually centered at RMSD>1Å with variable standard deviation. An example is the N-succinylamino acid racemases (M-CSA 377, EC: 4.2.1.113, reference PDB ID: **1R0M**), in which almost all of the catalytic residues exhibit some variation in location. This paradigm accounts for 33.9% of our enzyme families. (iii) **Two-state or open/closed** type: Here, the active site takes two distinct conformations, without evidence of intermediate states. Such behaviour appears in the RMSD histogram as a bimodal distribution, with highly variable mean and standard deviation of the peaks. An example is dihydrodipicolinate synthases (M-CSA 267, EC: 4.3.3.7, reference PDB ID: **1DHP**) where a very flexible, conserved and catalytically important Lys161 located on a loop, is captured in two distinct positions, with a transition possibly facilitating substrate binding[24]. Although this paradigm complies with the simplest interpretation of Koshland’s model of “induced fit”[8] (two perfectly distinct active site conformations, for the apo- and holo-forms respectively), it does not describe any more than 12.7% of our data. Interestingly, multi-state active sites are more common as described above. (iv) **Extensively variable**, a particularly interesting and rare case where a large fraction of the catalytic residues is captured in multiple different positions, suggesting a high degree of flexibility. The RMSD histogram here displays a non-regular shape and the catalytic residues are loosely clustered. An example of this behaviour is seen in protein-synthesizing GTPases (M-CSA 535, EC: 3.6.5.3, reference PDB ID: **1EFC**) where all catalytic residues are in loops or coils, in close contact to bulk solvent. This paradigm is the least frequent, including only 8.8% of the families. Last, there is a remaining 2.6% of families for which a classification could not be made as no conserved active sites were available.

### Case studies

In the following section we describe three distinct M-CSA homologous enzyme families which exhibit different types and degrees of flexibility in their catalytic sites. Fig. 5Fig. 7 present ensembles of structural analyses for each family studied respectively, illustrating active site flexibility by 3D superposition of conserved active sites, structural clustering to identify dominant conformers, as well as analyses such as secondary structure logos, and functional atoms RMSD distributions, both over the whole active site and for each residue. These examples were chosen to illustrate examples of flexible active sites and the relationship between the catalytic residue flexibility and the catalytic mechanism, and its biological relevance. The first example illustrates conservation of most of the active site, apart from one residue, which displays side chain flexibility; the second illustrates an enzyme with an open and closed state, in which two domains close up to form the active site (i.e., main chain and side chain flexibility); the third enzyme shows differential mobility between the catalytic residues, reflecting the type of ligand bound.

**Fig. 5:**
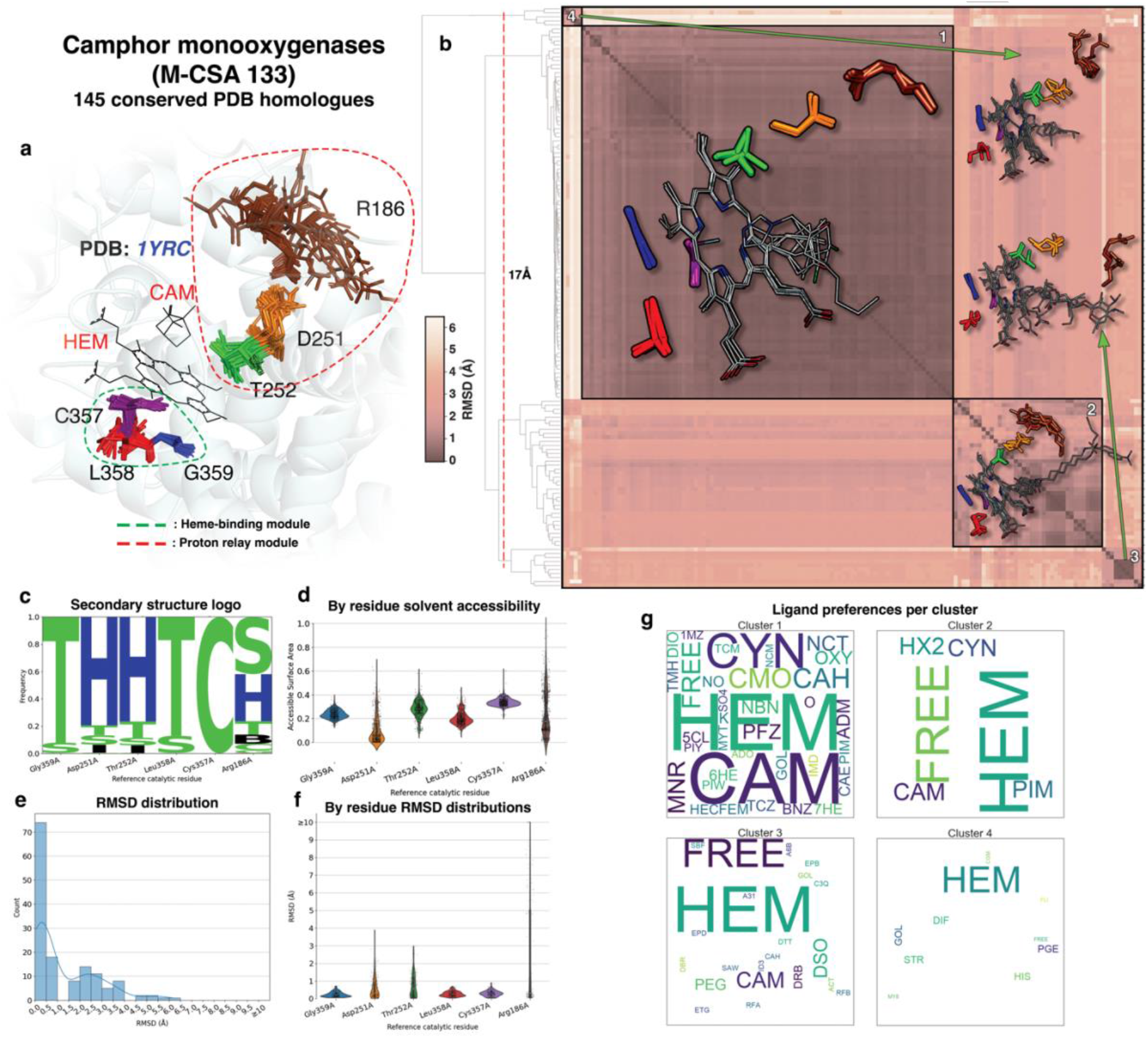
Active site structural analyses summary for P450cams. a: Conserved active sites from homologous enzymes are shown in superposition over the reference active site. Clusters of aligned catalytic residues are shown as distinctly coloured sticks and the structure of the reference enzyme of the family (PDB ID: **1YRC**) is shown in cartoon representation in the background. Ligands bound in the reference are shown in sticks and labeled with their PDB three-letter code. Residue groups forming individual modules are circled in dashed line. b: All-vs-all functional atoms RMSD matrix of conserved active sites. Darker and lighter colours, as shown in scale to the left of the matrix, indicate higher and lower structural similarity respectively. Clustering is performed by pruning the hierarchical dendrogram shown on the left of the matrix at a height indicated with a dashed line. Some representative members of each major cluster are overlayed on the matrix as sticks (side chain only) and lines (all atoms) for catalytic residues and bound ligands respectively. c: Secondary structure preferences for each residue position as assigned by DSSP (H: a-helix, E: β-strand, B: β-bridge, T: turn, S: bend, I: π-helix, C: random coil), presented in the form of frequency pseudo-sequence logo. d: Distributions of the Miller solvent accessibility for each residue position. e: Functional atoms RMSD for the whole active site. f: Functional atoms RMSD distributions for each residue. g: Word clouds of bound ligands (PDB three-letter codes) in each cluster of active sites. Size of words indicate frequency within the cluster. “FREE” word is used to indicate ligand-free active sites

#### Camphor 5-monooxygenases

Camphor 5-monooxygenases or P450cams (M-CSA 133, EC: 1.14.15.1, reference PDB ID: **1YRC**) are a well-studied family of cytochrome P450 enzymes that catalyse the hydroxylation of a camphor substrate, removing the requirement for a high temperature that would otherwise be essential without the enzyme[25]. P450cams are heme-bound enzymes and are known to change their conformation upon substrate binding[26]. Six catalytic residues constitute the active site of P450cam, forming two groups of distinct function (Fig. 5a): the first comprises consecutive residues Cys357, Leu358 and Gly359, responsible for heme binding, with Cys357 interacting with the heme Fe^2+^ ion (which has received an electron in a prior step by another protein, putidaredoxin)[27]. These three residues, as seen in the RMSD distributions of their functional atoms (Fig. 5c), remain very rigid within the family. The second triad of residues forms a proton relay responsible for transferring a proton from an oxidised peroxo moiety to bulk solvent. The transfer starts from Thr252 to a water molecule to Asp251, with Arg186 finally extracting it into bulk solvent. There is increasing “flexibility” of these three residues from the core to the surface of the protein, with Arg186 being extremely flexible and observed in either a conserved state, hydrogen bonded to Asp25, or highly variable location, when in contact with bulk solvent. Arg186 flipping is also reflected in the distribution of Miller solvent accessibility value (Fig. 5c), where two data-dense regions exist, corresponding to the two states of contact with bulk solvent.

Structural clustering of the active sites and enrichment analysis of each cluster according to the ligands bound (Fig. 5b and Fig. 5g respectively), show that Heme is bound to all structures. In the most populated and tight cluster (where Arg186 side chain is H-bonded to Asp251), camphor-like molecules are bound in the active site, whereas those active sites where Arg186 adopts a variable conformation, are either substrate-free or bind crystallographic artefacts and bulky molecules usually containing a covalently bonded camphor moiety (inhibitors or labelling agents). The overall conclusion here is that the major difference in the active sites lies in the conformation of Arg186, which is in a conserved, H-bonded state only when native-like camphor molecules are bound. Rigidity of the heme binding residues (on the other side of the heme) reflects the strong interactions with the heme, restraining them to a very defined position.

#### Peptidyl-dipeptidases (Angiotensin Converting Enzymes – ACEs)

Peptidyl-dipeptidases, usually known as Angiotensin Converting Enzymes (ACEs) (M-CSA 170, EC: 3.4.15.1, reference PDB ID: **1O8A**) catalyse the cleavage of various peptide substrates, including angiotensin I, enkephalins, kinins, amyloid peptides[28]. They are involved in blood pressure regulation in vertebrates by converting angiotensin I into its active form, angiotensin II, a vasoconstricting hormone (renin-angiotensin aldosterone system – RAAS)[29]. ACEs are highly promiscuous, accepting peptides of various lengths and physical-chemical properties, from which they usually cleave C-terminal dipeptides[29–32].

The catalytic site of these peptidases is large, with 8 residues (hydrogen bond donors and acceptors, electrostatic stabilisers and metal ligand binders), forming a groove between the N and C terminal lobes of the structure. Catalytic residues lie in both subdomains (Fig. 6a); substrate enters the groove between the two mobile lobes causing a transition from the open to the closed form. This flexibility allows the various substrates to enter and exit (“clam-shell”-like behaviour[28]). Catalytic activity depends on Zn^2+^ binding to a conserved His383-Glu384-Met385-Gly386-His387 (HExxH) motif in the reference ACE (PDB ID: **1O8A**). This motif is very well conserved in 3D (Fig. 6a), allowing us to define the Zn^2+^-binding 3D catalytic module as Hìs383_N_, Glu384_N_, His387_N_ and Glu411_N_ in the N-lobe (Glu411_N_ is not part of the HExxH motif, however it directly interacts with the ion). The second module is the group of His353_N_, Ala354_N_, His513_C_ and Tyr523_C_. The residue-by-residue RMSD distribution (Fig. 6e) shows that Zn^2+^ binding residues appear significantly more rigid compared to the rest. On the other hand, the three peptide-cleaving residues have higher RMSD values. Here, two types of flexibility can be described: (i) flexibility due to subdomain movement, where both main and side chain atoms vary significantly and equally (His513_C_ and Tyr523_C_) and (ii) side chain flipping, with main chain atoms remaining immobile (His353_N_). Interestingly, only two residues in the active sites are non-helical (His353_N_ and Ala354_N_ in Fig. 6c), but neither have a flexible backbone.

**Fig. 6:**
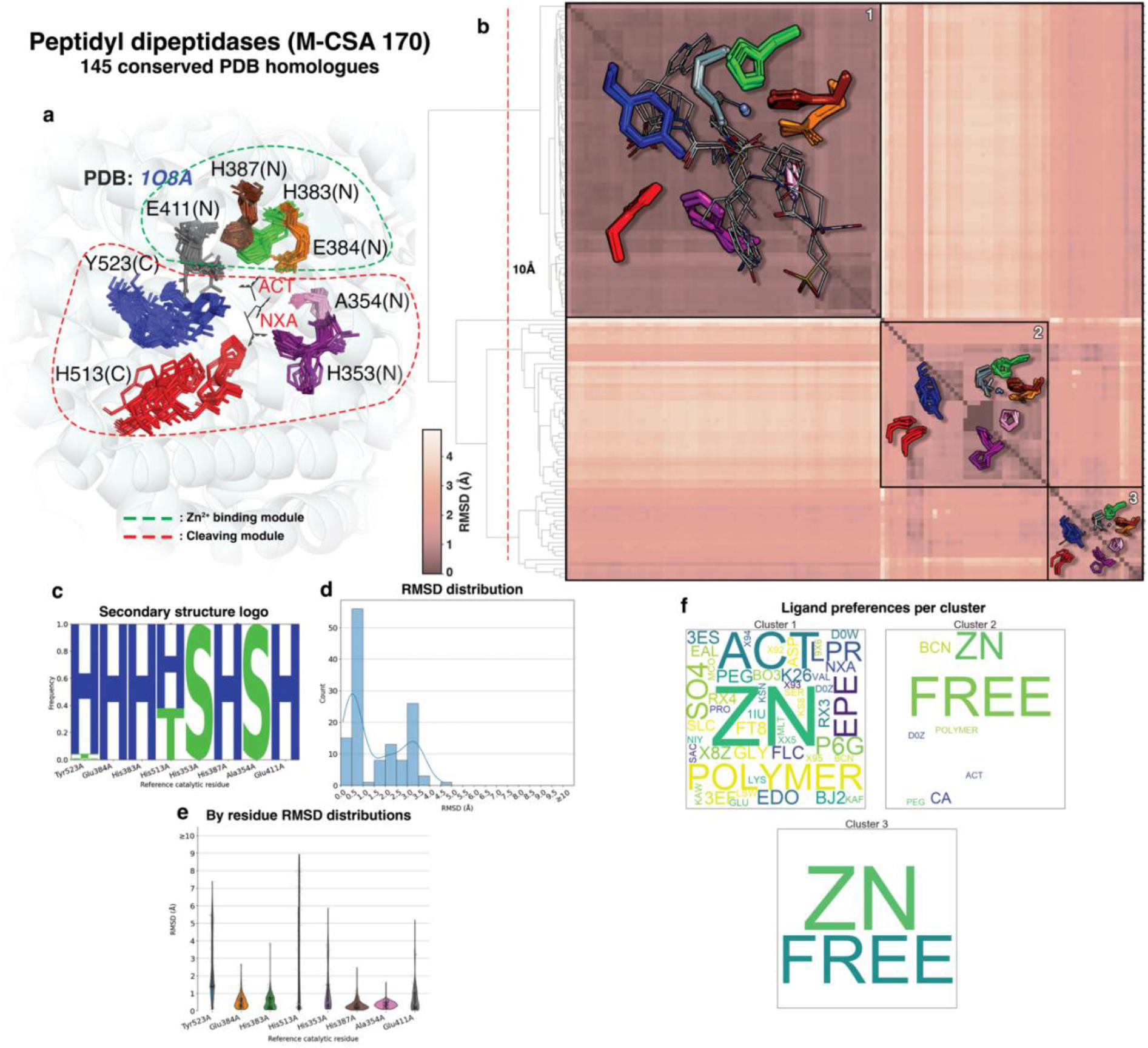
Active site structural analyses summary for Peptidyl dipeptidases. a: Conserved active sites from homologous enzymes are shown in superposition over the reference active site. Clusters of aligned catalytic residues are shown as distinctly coloured sticks and the structure of the reference enzyme of the family (PDB ID: **1O8A**) is shown in cartoon representation in the background (N and C letters in parentheses indicate the subdomain in which residues belong to). Ligands bound in the reference are shown in sticks and labeled with their PDB three-letter code. Residue groups forming individual modules are circled in dashed line. b: All-vs-all functional atoms RMSD matrix of conserved active sites. Darker and lighter colours, as shown in scale to the left of the matrix, indicate higher and lower structural similarity respectively. Clustering is performed by pruning the hierarchical dendrogram shown on the left of the matrix at a height indicated with a dashed line. Some representative members of each major cluster are overlayed on the matrix as sticks (side chain only) and lines (all atoms) for catalytic residues and bound ligands respectively. c: Secondary structure preferences for each residue position as assigned by DSSP (H: a-helix, E: β-strand, B: β-bridge, T: turn, S: bend, I: π-helix, C: random coil), presented in the form of frequency pseudo-sequence logo. d: Functional atoms RMSD for the whole active site. e: Functional atoms RMSD distributions for each residue. f: Word clouds of bound ligands (PDB three-letter codes) in each cluster of active sites. Size of words indicate frequency within the cluster. “FREE” and “POLYMER” words are used to indicate ligand-free sites and polymeric (protein or nucleic) bound ligands respectively.

**Fig. 7:**
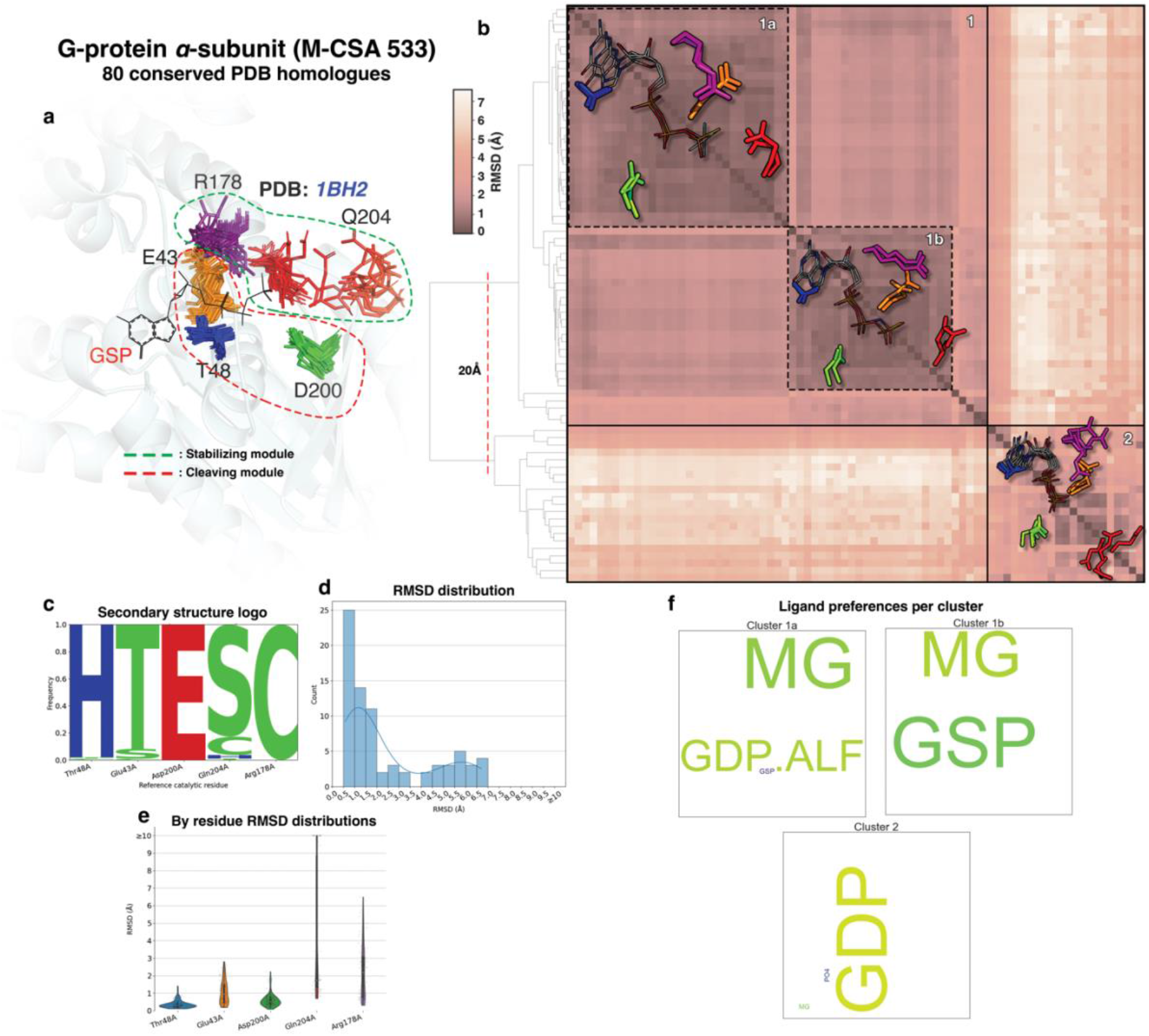
Active site structural analyses summary for the a-subunit of G-proteins. a: Conserved active sites from homologous enzymes are shown in superposition over the reference active site. Clusters of aligned catalytic residues are shown as distinctly coloured sticks and the structure of the reference enzyme of the family (PDB ID: **1BH2**) is shown in cartoon representation in the background. Ligands bound in the reference are shown in sticks and labeled with their PDB three-letter code. Residue groups forming individual modules are circled in dashed line. b: All-vs-all functional atoms RMSD matrix of conserved active sites. Darker and lighter colours, as shown in scale to the left of the matrix, indicate higher and lower structural similarity respectively. Clustering is performed by pruning the hierarchical dendrogram shown on the left of the matrix at a height indicated with a dashed line. Some representative members of each major cluster are overlayed on the matrix as sticks (side chain only) and lines (all atoms) for catalytic residues and bound ligands respectively. c: Secondary structure preferences for each residue position as assigned by DSSP (H: a-helix, E: β-strand, B: β-bridge, T: turn, S: bend, I: π-helix, C: random coil), presented in the form of frequency pseudo-sequence logo. d: Functional atoms RMSD for the whole active site. e: Functional atoms RMSD distributions for each residue. f: Word clouds of bound ligands (PDB three-letter codes) in each cluster of active sites. Size of words indicate frequency within the cluster.

Clustering the active site structures provides insight on ligand preferences (Fig. 6b and Fig. 6f). Three major clusters can be identified: one cluster of structures, whose conformation is well conserved (even for the relatively flexible His513_C_ and Tyr523_C_), all bind substrate or substrate-like molecules. The other two clusters exhibit significantly higher variance, and correspond to ligand-free sites or sites that bind low-MW crystallographic artefacts, like bicine (these are consistently located in the center of the catalytic module) (Fig. 6f). It should be mentioned here that both these clusters are dominated by ACE/SARS-CoV-2 spike protein complexes that are ligand free. For this enzyme family, binding of the substrate closes up the active site ready for catalysis; prior to ligand binding it is flexible and this surely provides the promiscuity it displays.

#### G-proteins, alpha subunit (GTPases)

G-proteins (M-CSA 533, EC: 3.6.5.-, reference PDB ID: **1BH2**) are membrane-bound cell signal transducers, propagating hormonal messages from the extracellular environment inside the cell, acting as switches for multiple downstream intracellular signalling cascades[33]. They are intrinsically slow-acting GTPases[34], in which GTP to GDP hydrolysis, and exchange of GDP for GTP by the G_α_ subunit, moderate the activity of the signal switch positively and negatively respectively, in a timed manner. A magnesium cofactor binds in the active site and there are five catalytic residues annotated in M-CSA, all acting as electrostatic stabilisers but with different specific roles[34,35] (Fig. 7a): Thr48, Asp200 and Glu43 are responsible for positioning and stabilising the substrate and a Mg^2++^ co-factor, constituting one of the two catalytic modules. Arg178 and Gln204 form the second module which contributes to the actual cleavage of γ-phosphate from GTP, passing through a dissociative transition state[35].

Structures of the alpha subunit of G-proteins with the PDB are usually crystallised in their holo-form, having bound either a slow-hydrolysable analogue of GTP (GTPγS) or a transition state analogue (GDP.AlF_3_)[36]. It is well known that significant conformational change occurs during catalysis in these enzymes, and different conformers are captured as snapshots in the crystal structures binding the analogues corresponding to the initial, intermediate, and final states of the reaction. The catalytically active module of Arg178 and Gln204 is affected the most, with both residues exhibiting high 3D variance as can be seen in the residue-by-residue RMSD distributions in Fig. 7e.

Structural clustering of the active sites (Fig. 7b and Fig. 7f) produces two major clusters, of which the first corresponds to GTPases binding either GTPγS or GDP.AlF_3_. This cluster is tight and includes two subclusters which mainly differ in the orientation of the Arg178 side chain. In the case of GTPγS (subcluster 1a), the Gln204 side chain points away from the γS phosphate group whereas when GDP.AlF_3_ (subcluster 1b) is bound the same side chain lies very close to the AlF_3_ moiety, demonstrating side chain and main chain flexibility. Similarly in the second major cluster, both Arg178 (Switch loop I) and Gln204 (Switch loop II)[37] are extremely variable, while the other residues remain relatively rigid. All active sites of the second cluster bind GDP (rather than a GTP analogue), with which Gln204 is unable to interact and is therefore flexible. The same applies to Arg178; this Arg finger, in agreement with the literature, passes through three states according to bound substrate[38]. Furthermore, minor but significant side chain flexibility can be seen in Asp200 which moves to coordinate the Mg^2++^ ion in a bidentate orientation between phosphate groups *β* and *γ* of the GTP analogue. Clearly these catalytic residues can transition among more than two conformational states during the progression of the catalytic mechanism, and residue-substrate interactions tend to stabilise a single conformer as clearly seen in the case of Gln204.

## Discussion

Our initial hypothesis, before doing the systematic analyses, was that most catalytic residues would exhibit 3D variation. This is only weakly supported by the results of this survey. Conversely, we find that enzyme families differ radically in their structural behaviour; some enzymes appear to be rigid; some enzymes exhibit significant flexibility in a subset of their catalytic residues, but the magnitude of flexibility and the number of catalytic residues that are flexible differs greatly between enzymes. Therefore, it is clear that no universal pattern of flexibility exists. We were able to derive a coarse qualitative classification of enzyme families according to the structural behaviour of their active site which indicates the pattern of dynamics of the catalytic toolkit and could provide a basis for more detailed quantitative classification. The main observation from this analysis is that the majority of enzyme catalytic sites tend to be rigid, with small, localised motions at the side chain level. The second most frequent behaviour is low-magnitude inherent flexibility (Fig. 4), where most residues tend to move slightly during catalysis. Although this paradigm is less frequent, it is of high significance and worth to be investigated systematically in the future. Last, all paradigms apart from “inherently rigid” are consistent with Koshland’s model of “induced fit”[8], (55.4% of enzyme families) with ligand binding being associated with the establishment of the biologically active conformation of the active site. Specifically, families of the “open/closed” paradigm (13%), adhere to the rudimentary form of this model, adopting two distinct conformational states, before and after ligand binding. Our results also demonstrate (especially in the case of G-proteins) that there can be a continuum of conformational states during catalysis in flexible enzymes. However, it is important to remember that these results reflect only those few snapshots of catalysis that are available in crystal structures. In these cases, transition state analogues give us insight of intermediate conformations a site might pass through, during the cycle of catalysis. A representative example supporting this, is the case G proteins (Fig. 7), as discussed further above. This work also aims to characterise the effect of ligands in enzyme active sites, with the term “ligand” being inclusive of substrates, co-factors and metals and exclusive of crystallographic artefacts. We demonstrate that active sites in their -apo form tend to be significantly more variable than in their -holo form, supporting the induced fit theory and the hypothesis that ligand binding leads to structural stabilisation of the enzyme. Moreover, binding of different ligands in the same active site will lead to different structural changes, as in the case of G-proteins (see above). Gutteridge and Thornton suggested that catalytic residue flexibility is usually subtle[11,12], based on a much smaller survey. This is in agreement with our results, which clearly show that the magnitude of functional group displacements is usually below 1Å, consistent with all the analyses in this paper. However, several examples of high order structural transitions do exist, with flexibility being accounted not only to rotameric shifts but also to whole-residue motion caused by domain or loop movement, as exemplified by the peptidyl-dipeptidase (Fig. 6) family where the flexibility of His513 and Tyr523 is attributed to opening and closing of the two subdomains of the protein before binding the peptide substrate.

We also explored whether catalytic residue flexibility reflects the “catalytic role” of the residues[2]. We found that although huge variation among enzyme families exists and the degree of correlation varies, there are some consistent preferences: “spectator” residues are more flexible than “reactant” residues in general, metal binding residues are rigid, while charged residues tend to be more flexible. Again, although these patterns do exist, they are not absolute and vary among different enzyme families. Generally, residues directly involved in the chemistry of catalysis (like nucleophiles and proton donors/acceptors) are rigid, with 3D variation lying within experimental error limits. A caveat should be stated here: This work has concentrated only on catalytic residues; hence further investigation is required to provide more light on whether similar trends also occur for residues in the binding site or residues not involved in catalysis at all. Furthermore, association of active site 3D variation/flexibility with enzyme kinetics is a topic we have yet to explore.

Enzymes utilise a limited toolkit of residues and co-factors to perform a vast number of different catalytic reactions[2,39], with many enzymes being promiscuous and more general-purpose than others. Exploring the various catalytic mechanisms curated with M-CSA, one could perceive a common trait: in many cases the catalytic residues can be grouped according to the function they perform during the progression of catalysis. In this context, we introduce the concept of “functional module”: a self-contained cluster of catalytic residues participating in a well-defined part of the catalytic mechanism (e.g., the catalytic triad or a Zn^2++^-binding cluster of histidines). The case studies reported here show that residues of the same module tend to loosely follow a similar structural transition trend, however exceptions do exist, and this further complicates the definition of the term. This initial description will be enhanced in the future, to aid in better understanding catalysis and designing enzymes of novel function.

The scope of this paper is to examine the phenomenon of 3D variation within identical or similar enzymes, without exploring plasticity of active sites[6] and convergent evolution[40]. However, the computational pipeline implemented during this work (**CSA-3D**) is designed to allow the study of plasticity, by generating active site 3D templates[41–44]. Template generation is straightforward for rigid active sites; however, sites of high flexibility cannot be easily represented in one template. However, since **CSA-3D** supports clustering of the structures, it is now possible to generate templates corresponding to alternative conformers of the same active site. Templates can be used to query protein structures to identify catalytic entities, making them a robust tool (and alternative to sequence alignment) to explore cases of functional promiscuity, moonlighting and convergent evolution and to aid the design of novel enzymes. Source code of **CSA-3D** is freely available at *https://github.com/iriziotis/CSA-3D*. Although current version of **CSA-3D** is purpose-made for the analyses presented in this paper, we plan to eventually release it in the form of a public tool to generate enzyme structural reports, and also include these reports (similar to the case studies presented further above) in M-CSA, covering all families.

## Methods

### Catalytic residue annotations

Catalytic residue information for a total of 925 homologous enzyme families was obtained from the Mechanism and Catalytic Site Atlas (M-CSA)[14] using the public API. Each entry in M-CSA consists of a reference enzyme structure in which catalytic residues are manually annotated, and a set of sequence homologues from the Protein Data Bank[45] identified by PHMMER[46], where catalytic residues are annotated through multiple sequence alignment. Homology is defined by an E-value threshold of 10^−6^. Annotations include basic information obtained from PDBe[47] (PDB ID, residue name and residue identifer) as well as curated information related to the function of each catalytic residue (functional role in reaction, post-translational modification, and location of function (main chain or side chain)).

### Catalytic residue 3D coordinates

3-dimensional coordinates of catalytic residues were extracted from PDBe biological assembly structures (total number of structures covering the whole M-CSA: 62,343). The sequence-based method used to find homologues within the PDB identifies, for each reference catalytic residue, the residue name and its position in the sequence of all homologous chains in the PDB file. Sometimes active sites are composed of residues from different subunits and many enzymes contain multiple copies of an active site. Therefore, it is necessary to explicitly identify the chain or chains that contribute catalytic residues to each active site.

### Reconstruction of active sites in 3D

A downstream processing step was applied to reconstruct clusters of catalytic residues constituting all active sites in an “assembly enzyme structure”. We used the following clustering algorithm to identify all active sites in each assembly: a random catalytic residue from a structure is selected and all the residues sharing the same name and position in the chain sequence (equivalent residues located in other chains) are set as the seed of a new cluster. Each cluster corresponds to one catalytic site; site reconstruction is performed by iterating through all catalytic residues in all chains and adding the one closest to the center of mass of the current catalytic site (cluster). An exception had to be addressed regarding symmetric active sites, such as those of HIV proteases, where the active site lies in the interface of two identical subunits[48] and contains pairs of equivalent residues. To account for these unique cases, an extra identifier was assigned to each curated catalytic residue in M-CSA and the equivalence criterion included this identifier along with the residue name and residue identifier, so that ambiguity is eliminated.

The structural “sanity” of the reconstructed active sites also had to be checked; non-standard residue and chain numbering in some PDB structures might cause the construction of irregular and biologically non-meaningful active sites. All sites were checked by comparing all the inter-residue distances with the corresponding distances of the reference catalytic site (which is considered the ground truth since the chain identifier of its catalytic residues is manually annotated) and of all other catalytic sites extracted from the same structure. The sanity criteria are the following: All inter-residue pairwise distances above 8Å must be 3 times or less their corresponding distances in the reference active site. A similar criterion is applied when comparing to sites of the same structure, but in this case the cut-off is 1.3 times the distance of the equivalent residue pair. Active sites not satisfying these criteria were filtered out and the rest passed through another filter of redundancy reduction. If a structure contains multiple structurally similar active sites, with an RMSD threshold of 0.5Å, only one instance of the active sites is kept^6^. This is also done to eliminate redundant active sites originating from copying chains of the asymmetric unit into the biological assembly. The distance thresholds in the filtering process are arbitrary and were defined empirically by observing several cases of incorrect active site construction. The RMSD threshold we used to reduce redundancy within the same structure was also empirical and corresponds to an estimate of the discrepancy of atomic positions due to resolution experimental error.

### Ligand identification and annotation

The next step in active site construction was the identification of nearby ligands and their classification as substrates, co-factors, or artefacts. Ligands adjacent to the catalytic residues are identified by a k-d tree nearest neighbor search[49] algorithm implemented in the BioPython package[50]. First, a spherical search volume of 3Å radius around each atom of the catalytic residues is defined using its 3D atomic coordinates, and all non-protein components and polymer residues not belonging to the chains constituting the active site are captured as potential ligands interacting with the active site. It should be noted that polymer ligands (e.g., in proteases or DNA-interacting enzymes) are treated as a single segment and not as individual residues. Moreover, empty space not covered by the search volume (e.g., between distant catalytic residues whose search “spheres” do not overlap (i.e. 6Å maximum distance between the centres of search)) is filled by adding pseudo-atoms (search centres) positioned in the gap between the distant residues. The exact coordinates of the interpolated search centers are calculated as the average center-of-mass of the two residues between which the empty space must be filled. In the second step, non-protein co-factor-like and substrate-like components located peripherally to the active site are also captured and annotated as “distal” by searching a volume of 30Å radius, centered at the active site centroid.

Classification of ligands in categories such as co-factors, substrates and artefacts is performed in the next step, by applying the following pipeline for each ligand: First the chemical similarity of the ligand with all the cognate reactants and products of the native enzyme reaction is calculated. This is done by mapping the protein structure to an enzyme reaction via its E.C. number as collected from SIFTS[51], obtaining the corresponding reactant and product chemical information from M-CSA, Rhea[52] or KEGG[53] and calculating the PARITY (<u>P</u>roportions of <u>A</u>toms <u>R</u>esiding in <u>I</u>dentical <u>T</u>opolog<u>Y</u>) chemical similarity score[19] between the ligand and each component. The ligand is then annotated, using the match with the highest score, with its corresponding cognate component. It is important to note that PARITY score cannot be computed for complex polymers and single-atom components thus no corresponding match is linked to these ligands. Second, ionic, and non-ionic co-factors as well as other single-atom components (metals, non-metals, noble gases) are annotated using a PDB identifier to three-letter code mapping kindly provided by PDBe (which collects information from the Co-Factor Database[54]). Third, components known to be used to facilitate protein crystallisation are annotated as artefacts if they do not satisfy a PARITY score of >30% with at least one cognate reactant, product, or co-factor. Last, to capture the location of the ligand relative to the “active site”, each ligand is annotated with a metric of centrality relative to the catalytic residues; this is the average center-of-mass distance of the ligand to the catalytic residues, normalised over the average center-of-mass inter-residue distance^7^.

### Superposition of homologous active sites

Structural comparison of active sites was performed by structural superposition over a subset of three functional atoms per catalytic residue[43,55]. Atom triads are selected based on the function location of the catalytic residues as annotated in M-CSA; for instance, for an Arg residue functioning via its side chain, the superposition atoms will be the three endpoint guanidino atoms of its side chain (CZ, NH1, NH2 as defined in the PDB format). Likewise, for residues that function via their main chain, or are subject to post-translational modifications, three main chain atoms are selected. The exact definitions are presented in Fig. S1. This approach allows for precise superposition on the functionally important segment of the residue, allowing for the non-functional atoms to be more variable in the 3D space, without biasing the alignment.

Catalytic residue mutations are also handled during superposition. We define three types of residue mutations: the first type are mismatches between two residues of similar physical chemical properties (Asp ↔ Glu, Asn ↔ Gln, Ser ↔ Thr ↔ Tyr, Val ↔ Leu ↔ Ile); these cases are treated by selecting common functional atoms between the residues. The second type are mismatches between non-equivalent residues; in this case, pseudo-mutations on the functional atoms are introduced (in the form of a common atom identifier indicating a mismatch), to facilitate alignment between atoms of the same type. Finally, the third type regards non-aligned residues (in sequence) which are defined as gaps; here, the residue/gap pair is completely excluded from superposition. It is also important to take into account the symmetry of functional groups in some residues, i.e. the two terminal O atoms of the Asp and Glu carbonyl group. These groups might appear flipped in two similar crystal structures solved in different experiments, due to ambiguity when assigning atoms on the electron density map. A special case is the His imidazole ring, in which the N and C atoms cannot be easily distinguished by their density clouds alone, therefore these atoms must be treated as equal, allowing for flipping of the side chain[56]. This problem is addressed by inserting wildcards to the identifier of ambiguous functional atoms and applying an assignment algorithm to find the optimal one-to-one atom correspondence. We used the Kuhn-Munkres algorithm, also known as the Hungarian algorithm on the ambiguous functional group pairs, after an initial crude Kabsch superposition[57] of the active sites, which solves the assignment problem by considering the atom identifier wildcards and their pairwise Euclidean distances on the 3D space[58]. In non-equivalent residue mismatches, all functional atoms are treated as equal, and the same algorithm is applied to find the optimal correspondence.

Fitting via RMSD minimisation, by definition, will find the superposition with the lowest global RMSD; for example, in a 6-residue active site, where 5 are rigid and 1 is highly mobile, the RMSD will be shifted to compensate the deviation of this one residue – creating an average which does not reflect the rigidity of the core 5 and flexibility of the outlying residue. Therefore, to account for this partial structural variability in active sites, fitting had to be performed in a dynamic manner, applying positive bias to invariable regions of the active site, and negative bias to more unstable regions (e.g., residues of a flexible “lid” loop in Histidine kinases[59]). The method applied to facilitate this is a Gaussian-weighted RMSD superposition[60], a variant of the classical Kabsch algorithm, specially implemented for our purposes. A detailed mathematical description of the algorithm can be found in the Supplementary Material document.

### Overall sequence and structural comparison of homologous enzymes

Comparison of enzyme sequences was performed by pairwise sequence alignment using the BioPython package[61], while structural comparison at the overall structure level was done by running *TM*-align[62] on the C_α_ atoms of each chain. To calculate these parameters in enzymes composed of multiple subunits, homologous chains that form the active site are superimposed separately, and the final metric is averaged over the number of chain pairs compared.

### Data clean-up

For sequence vs. structure analyses, 30,859 reference/homologue comparisons were used, deriving from 761 homologous enzyme families after applying a resolution filter of 2.0Å and excluding NMR structures. To reduce bias by over-representation of heavily populated families (e.g. members of kinase and protease superfamilies), the number of comparisons per family was capped at 30, and a random sample was collected. Additionally, a maximum RMSD filter of 10Å (overall structure and active site C_α_ atoms) was applied to the dataset, to reduce the presence of artefacts. This RMSD threshold was selected empirically after inspection of individual comparisons to minimise the presence of artefacts, with minimal exclusion of biologically relevant examples. The final dataset for this set of analyses covers 10,430 unique PDB structures.

For analyses of the effect of ligands in active sites, again only non-NMR structures of ≥2.0Å were considered. Furthermore, enzymes with no EC number assigned and active sites containing gaps (non-aligned catalytic residues) were excluded. It is expected that active sites of the same family will display similar clustering, as expressed by the mean intra-residue distance of an active site. Assuming that an extreme mean intra-residue distance discrepancy (>5Å) between two homologous sites is probably a result of erroneous site building, pairs like this are considered artefacts and are excluded. RMSD filters were also applied as follows: pairs with wRMSD>10Å or maximum single residue RMSD>20Å were excluded, as well as pairs of wRMSD>1.5Å where maximum single residue RMSD is more than 3 times greater than the wRMSD value (see superposition algorithm description in Supplementary Material for wRMSD definition). Finally, the dataset comprised 2,793,916 comparisons from 796 M-CSA families, covering 21,039 unique PDB structures. Similar data filtering was applied for the analyses of the effect of residue types and roles on 3D variability.

### Structural analyses within homologous active sites

A series of analyses was performed on all homologous enzyme families in M-CSA for which active site 3D information could be adequately extracted. The first two are secondary structure assignment and solvent accessibility of each catalytic residue, both calculated by DSSP[63], with solvent accessibility being expressed with the Miller accessible surface area metric[64].

The next calculation was to estimate the structural variation among homologous active sites. Initial screening of conformational heterogeneity was performed by all-vs.-all active site superposition and superposition of the reference enzyme active site with all homologous sites. *RMSD* values were computed via the weighted superposition method described in the previous paragraph, aligning over functional atom triads for each catalytic residue. Additionally, to view the structural variability each catalytic residue might exhibit, *RMSD* was measured separately on each residue, again on an all-vs.-all and reference-vs.-all basis.

Last, all-vs.-all RMSD values were put into a dissimilarity matrix, which was used to cluster the active structures. Clustering was performed by linkage analysis on the RMSD matrix and construction of a hierarchical dendrogram[21]. Clusters were derived by pruning the dendrogram at an empirical RMSD height equal to 0.4 times the maximum RMSD value of the linkage matrix.

## List of PDB IDs

1NIA,1R0M,1DHP,1EFC,1YRC,1O8A,1BH2

## Author contributions

IGR: Conceptualisation, Software, Methodology, Validation, Formal analysis, Investigation, Visualisation, Writing – Original Draft preparation

AJMR: Data Curation, Writing – Review & Editing

NB: Validation, Writing – Review & Editing

JMT: Conceptualisation, Supervision, Resources, Funding Acquisition, Project Administration, Writing – Review & Editing

## Acknowledgements

The work was supported by the EMBL International PhD Programme (IGR) and the European Molecular Biology Laboratory (AJMR, NB, JMT)

