## Supplementary Information for "Conformational variation in enzyme catalysis: A structural study on catalytic residues"

### Supplementary Material

##### Active sites dynamic superposition algorithm

For two homologous active sites *p* and *q*:

Functional atoms 3D coordinate sets $P$ and $Q$ respectively, of equal dimensions $N \times 3$, are extracted, where $N$ is the number of atoms. Coordinates $Q$ are initially superposed to $P$, applying the Kabsch algorithm in a single step, followed by calculation of unweighted *RMSD*

$$RMSD = \frac{1}{N}\sqrt{\sum_{i=1}^{N} d_i^2}$$

where $d_i$ is the pairwise Euclidean distance of two corresponding atoms. This initial RMSD value is used to compute a scaling factor *c* used downstream in the calculation of atom weights. The scaling factor is calculated by a sigmoid function, defined empirically, after examining the superposition of active sites within several homologous enzyme families

$$c = \frac{9}{1 + e^{(-1.7RMSD-5.3)}} + 1$$

Atom weights (as vector *W*) are derived by incorporating the pairwise atom Euclidean distances into a Gaussian function, scaled by the scaling factor from the previous step, as

$$W = (w_{i=1}, \quad \cdots \quad, w_{i=N})$$

where

$$w_i = e^{-\frac{d_i^2}{c}}$$

Next, the centroids of the two coordinate sets are calculated as

$$C_P = \begin{pmatrix} \frac{1}{N}\sum_{i=1}^{N} w_i x_{Pi} \\ \frac{1}{N}\sum_{i=1}^{N} w_i y_{Pi} \\ \frac{1}{N}\sum_{i=1}^{N} w_i z_{Pi} \end{pmatrix}, C_Q = \begin{pmatrix} \frac{1}{N}\sum_{i=1}^{N} w_i x_{Qi} \\ \frac{1}{N}\sum_{i=1}^{N} w_i y_{Qi} \\ \frac{1}{N}\sum_{i=1}^{N} w_i z_{Qi} \end{pmatrix}$$

Coordinates are then translated to the origin of the cartesian space by subtracting the weighted centroids

$$P' = P - C_P$$
$$Q' = Q - C_Q$$

The next step is the calculation of a weighted cross-covariance matrix between the two coordinate sets

$$K_{P'Q'} = (P'^T W)Q'$$

The cross-covariance matrix is factorised through singular value decomposition (SVD)

$$K_{P'Q'} = U\Sigma V^T$$

Rotation matrix is calculated as

$$R = (VU^T)^T$$

Translation vector is calculated as

$$T = C_Q - C_P R$$

$C_Q$ coordinates are superposed on the $C_P$ coordinates by applying a transformation operation using the rotation matrix *R* and translation vector *T*

$$Q'' = Q'R + T$$

Finally, the same atom weights are incorporated into *RMSD* calculation (weighted *RMSD*, or *wRMSD*)

$$wRMSD = \frac{1}{N}\sqrt{\sum_{i=1}^{N} w_i d_i^2}$$

The process is repeated by calculating new weights, weighted transformation and *wRMSD* until a *wRMSD* convergence of < 0.0001Å is achieved, with a maximum of 6000 iterations. In a few cases where convergence cannot be achieved, unweighted superposition is performed in 10 cycles, applying simple rejection of outlying atoms, with a pairwise distance threshold of 5Å. After successful superposition of the homologous active sites, a final, unweighted *RMSD* value over all functional atom triads is calculated. This approach not only accomplishes optimal, flexibility-aware superposition, but also outputs an *RMSD* value proportional to the overall structural dissimilarity of the active sites. The reason why unweighted *RMSD* is more appropriate for our purposes is because *wRMSD,* in principle, reflects the dissimilarity of only the most rigid parts of a structure, weighting down the more variable parts, therefore it is not informative of the overall dissimilarity.

### Figures

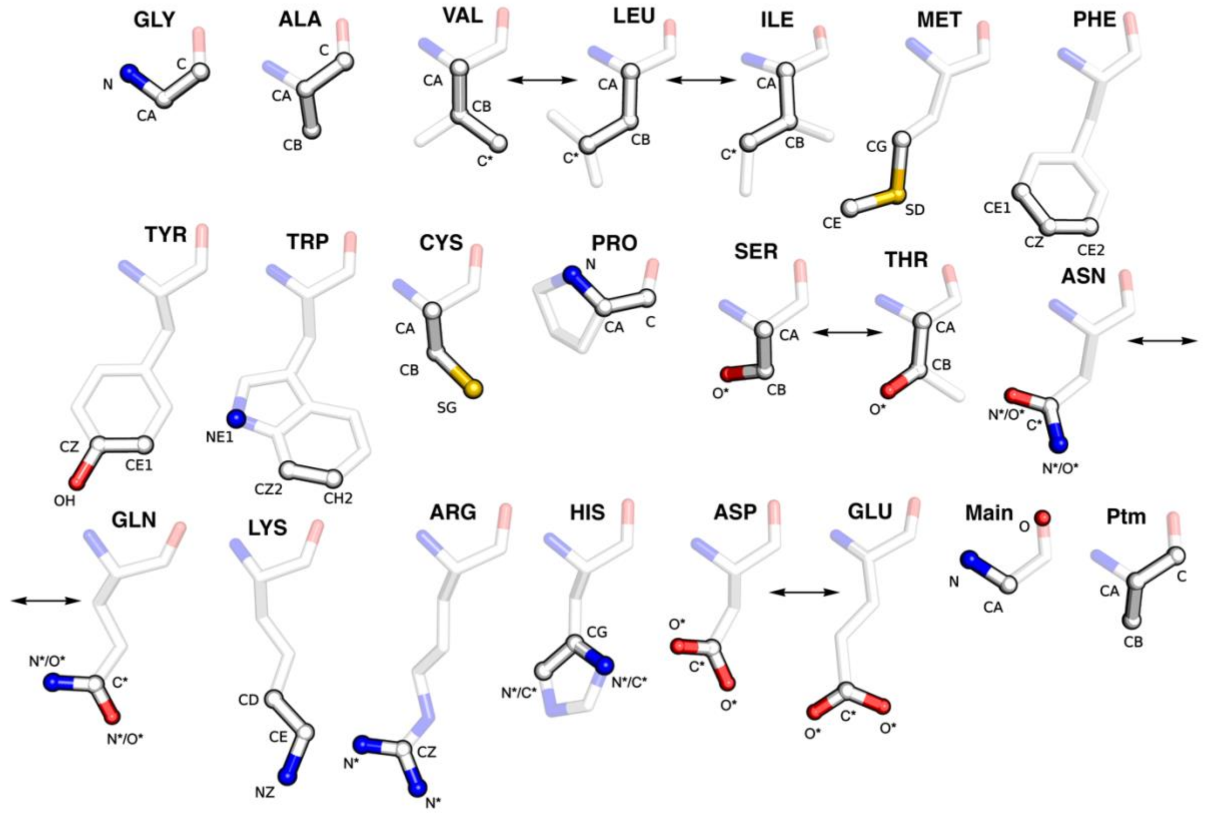


*Fig. S1: Functional atoms used in superposition of homologous active sites. For each residue type, three residue-specific atoms are selected (shown in opaque sticks) if the residue functions via its side chain. In cases where a residue functions via its main chain or undergoes post-translational modifications, explicit atom selections are defined, as shown in the lower right part of the figure. Bidirectional arrows indicate residues of equivalent properties that can be superposed interchangeably. Similarly, atoms of symmetrical chemical groups or atoms that are shared between equivalent residues are indicated with a * symbol. These atom selections are a modified version of the definitions from Wallace et al*[1]. *Models were generated in PyMol*[2].

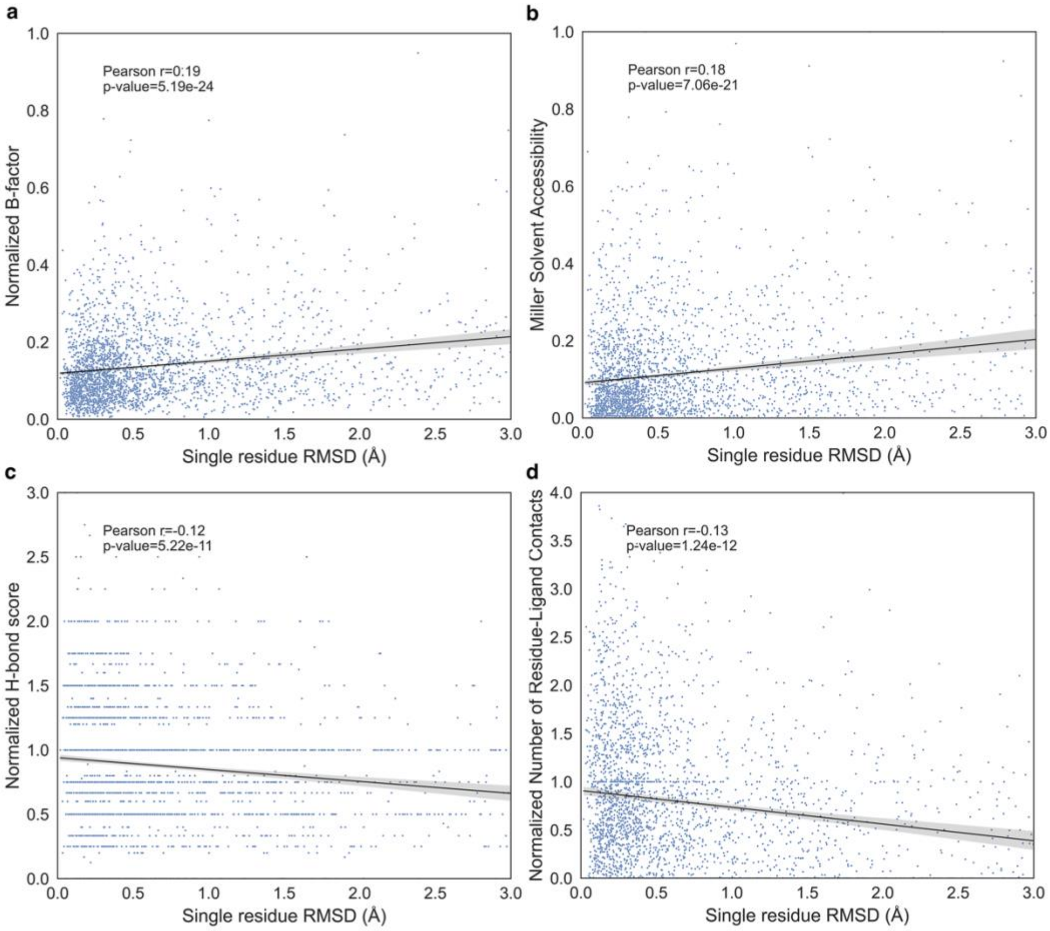


*Fig. S1: Correlation between single-residue RMSD versus various structural parameters. a: Normalized B-factor (normalization performed over the Z-score of all B-factors of the structure), b: Miller Solvent Accessibility, c: Normalized H-bond score (expressed as the number of putative polar contacts of donor/acceptor atoms of a residue with neighbouring molecules using a distance threshold of 3.5Å, normalized over the total number of its donor/acceptor atoms), d: Normalized number of residue-ligand contacts (expressed as the number of contacts between atoms of a residue and neighbouring ligands, using a distance threshold of 8Å, normalized of the total number of residue/ligand atoms). Results derive from superposition of active sites within homologous M-CSA families with their respective reference active site. Linear regression was applied in each case, indicated as a line, while the corresponding Pearson correlation coefficients (r) and their p-values are annotated.*

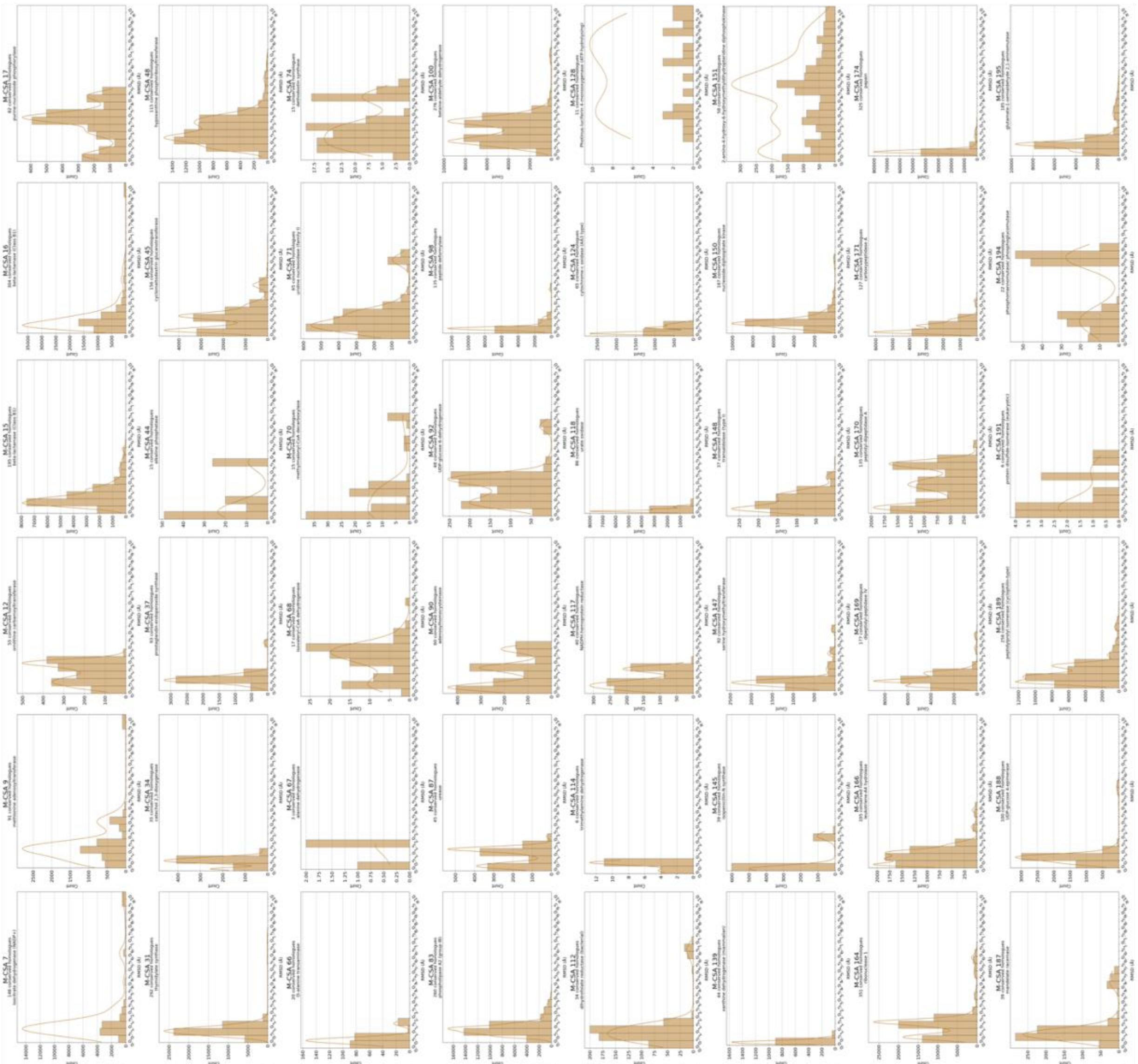

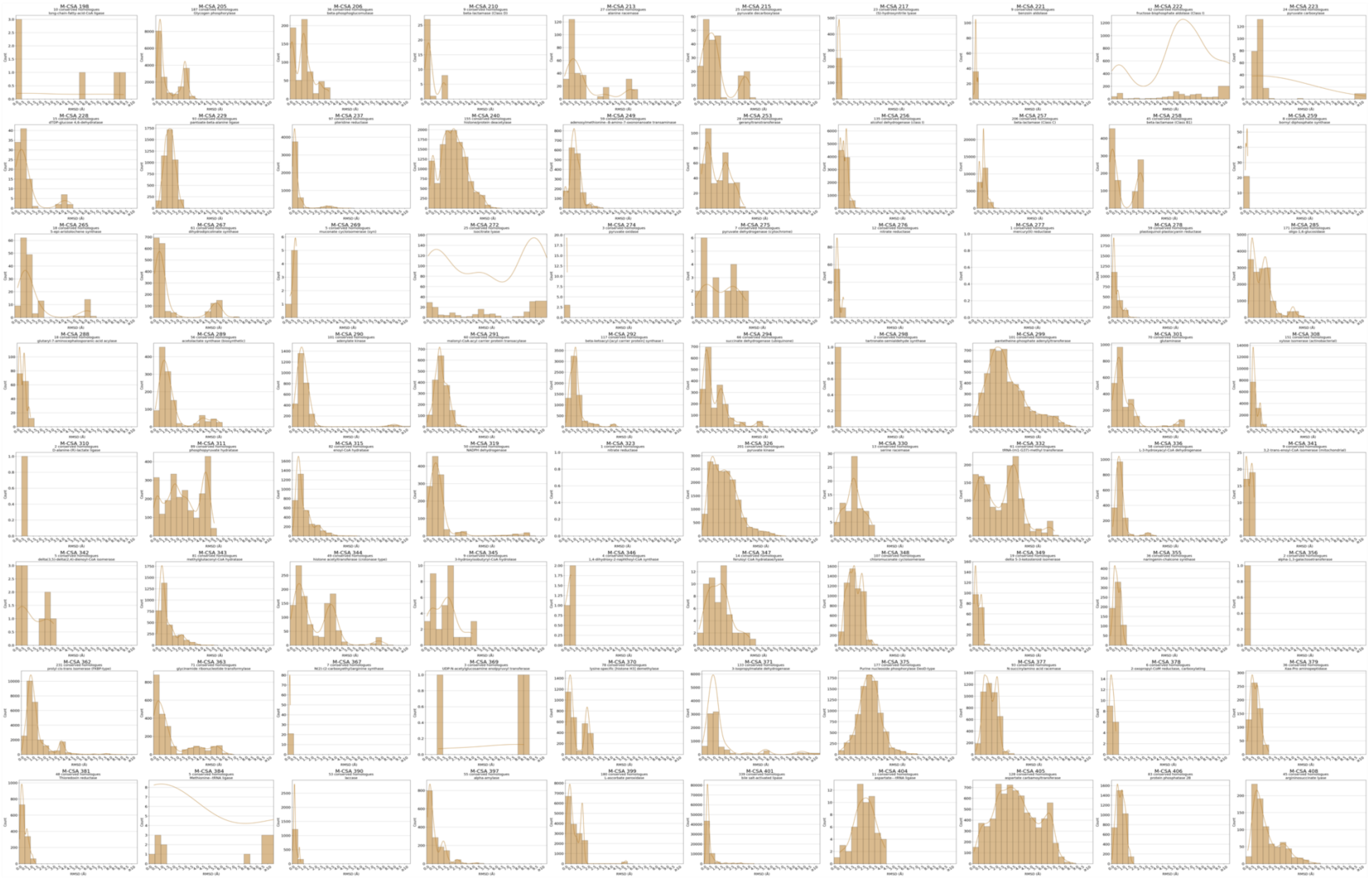
M-CSA 198
M-CSA 205
M-CSA 206
M-CSA 210
M-CSA 213
M-CSA 215
M-CSA 217
M-CSA 221
M-CSA 222
M-CSA 223
M-CSA 228
M-CSA 229
M-CSA 237
M-CSA 240
M-CSA 249
M-CSA 253
M-CSA 256
M-CSA 257
M-CSA 258
M-CSA 259
M-CSA 265
M-CSA 267
M-CSA 269
M-CSA 272
M-CSA 274
M-CSA 275
M-CSA 276
M-CSA 277
M-CSA 278
M-CSA 285
M-CSA 288
M-CSA 289
M-CSA 290
M-CSA 291
M-CSA 292
M-CSA 294
M-CSA 298
M-CSA 299
M-CSA 301
M-CSA 308
M-CSA 310
M-CSA 311
M-CSA 315
M-CSA 319
M-CSA 323
M-CSA 326
M-CSA 330
M-CSA 332
M-CSA 336
M-CSA 341
M-CSA 342
M-CSA 343
M-CSA 344
M-CSA 345
M-CSA 346
M-CSA 347
M-CSA 348
M-CSA 349
M-CSA 355
M-CSA 356
M-CSA 362
M-CSA 363
M-CSA 367
M-CSA 369
M-CSA 370
M-CSA 371
M-CSA 375
M-CSA 377
M-CSA 378
M-CSA 379
M-CSA 381
M-CSA 384
M-CSA 390
M-CSA 397
M-CSA 399
M-CSA 401
M-CSA 404
M-CSA 405
M-CSA 406
M-CSA 408
Count
RMSD (Å)

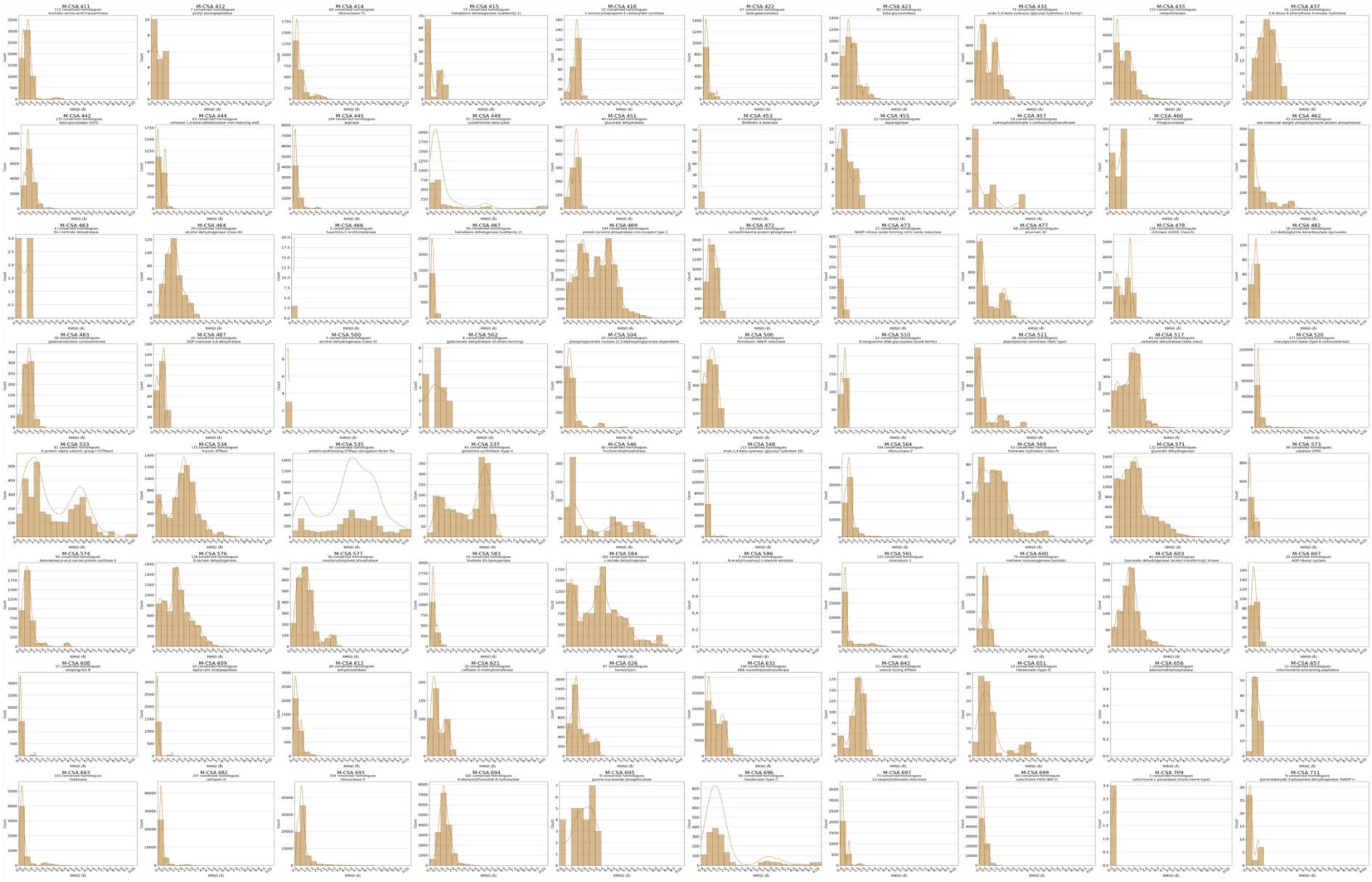
M-CSA 411
M-CSA 412
M-CSA 414
M-CSA 415
M-CSA 418
M-CSA 422
M-CSA 423
M-CSA 432
M-CSA 433
M-CSA 437
M-CSA 442
M-CSA 444
M-CSA 445
M-CSA 449
M-CSA 451
M-CSA 453
M-CSA 455
M-CSA 457
M-CSA 460
M-CSA 462
M-CSA 463
M-CSA 464
M-CSA 466
M-CSA 467
M-CSA 469
M-CSA 472
M-CSA 473
M-CSA 477
M-CSA 478
M-CSA 482
M-CSA 483
M-CSA 487
M-CSA 500
M-CSA 502
M-CSA 504
M-CSA 506
M-CSA 510
M-CSA 511
M-CSA 517
M-CSA 520
M-CSA 533
M-CSA 534
M-CSA 535
M-CSA 537
M-CSA 546
M-CSA 548
M-CSA 564
M-CSA 569
M-CSA 571
M-CSA 573
M-CSA 574
M-CSA 576
M-CSA 577
M-CSA 583
M-CSA 584
M-CSA 586
M-CSA 591
M-CSA 600
M-CSA 603
M-CSA 607
M-CSA 608
M-CSA 609
M-CSA 612
M-CSA 621
M-CSA 626
M-CSA 632
M-CSA 642
M-CSA 651
M-CSA 656
M-CSA 657
M-CSA 663
M-CSA 682
M-CSA 693
M-CSA 694
M-CSA 695
M-CSA 696
M-CSA 697
M-CSA 699
M-CSA 709
M-CSA 711

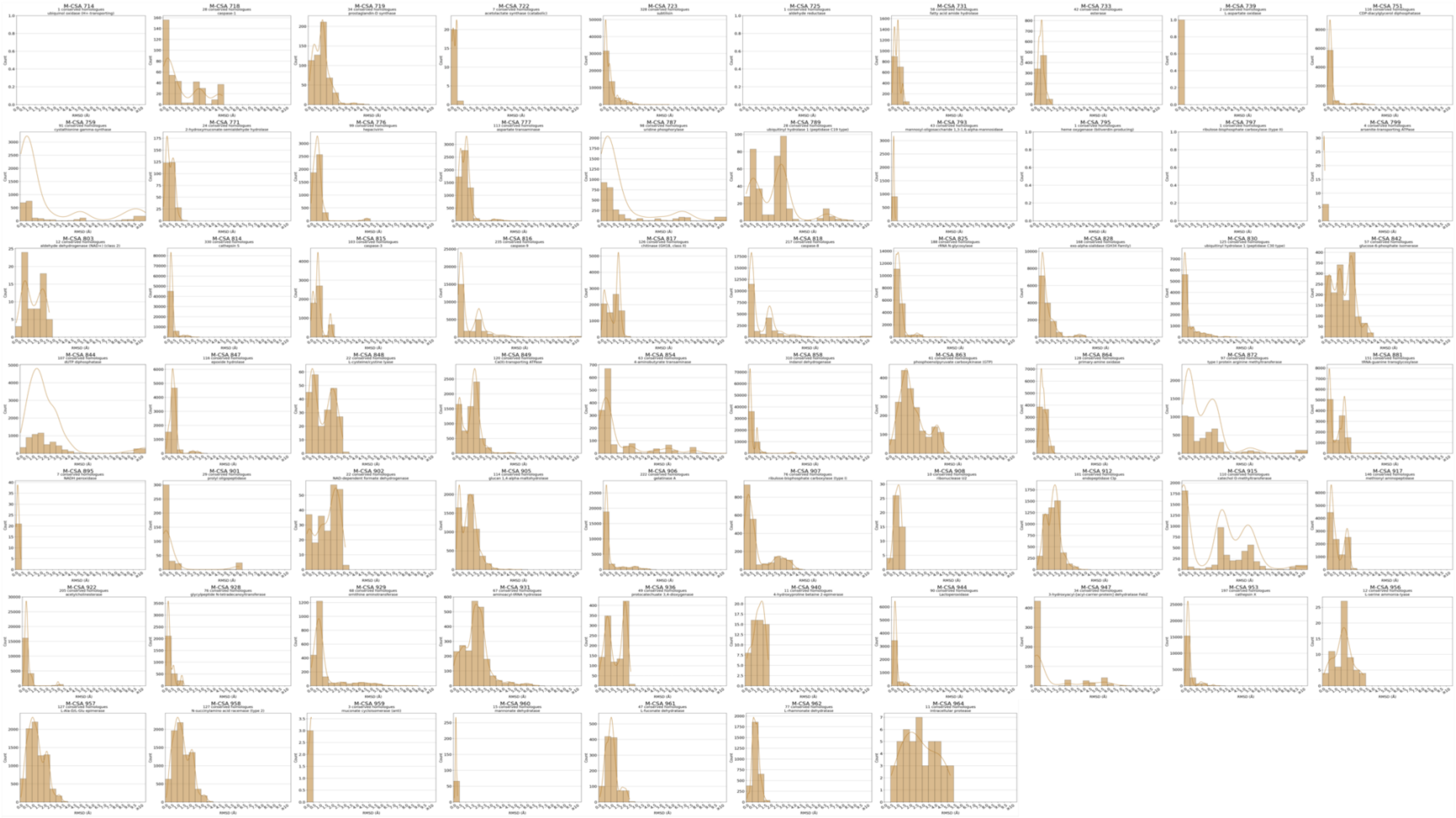
M-CSA 714
M-CSA 718
M-CSA 719
M-CSA 722
M-CSA 723
M-CSA 725
M-CSA 731
M-CSA 733
M-CSA 739
M-CSA 751
M-CSA 759
M-CSA 771
M-CSA 776
M-CSA 777
M-CSA 787
M-CSA 789
M-CSA 793
M-CSA 795
M-CSA 797
M-CSA 799
M-CSA 803
M-CSA 814
M-CSA 815
M-CSA 816
M-CSA 817
M-CSA 818
M-CSA 825
M-CSA 828
M-CSA 830
M-CSA 842
M-CSA 844
M-CSA 847
M-CSA 848
M-CSA 849
M-CSA 854
M-CSA 858
M-CSA 863
M-CSA 864
M-CSA 872
M-CSA 881
M-CSA 895
M-CSA 901
M-CSA 902
M-CSA 905
M-CSA 906
M-CSA 907
M-CSA 908
M-CSA 912
M-CSA 915
M-CSA 917
M-CSA 922
M-CSA 928
M-CSA 929
M-CSA 931
M-CSA 936
M-CSA 940
M-CSA 944
M-CSA 947
M-CSA 953
M-CSA 956
M-CSA 957
M-CSA 958
M-CSA 959
M-CSA 960
M-CSA 961
M-CSA 962
M-CSA 964
RMSD (Å)
Count

*Fig. S2: All-vs-All functional atoms RMSD distributions of conserved active sites, for all M-CSA entries containing at least 50 homologous active site structures. M-CSA ID, enzyme name and number of conserved active sites are annotated on each histogram plot. RMSD scale (x-axis) has a common range for all distributions. RMSD values of ≥10Å are binned altogether for better visualization. Bar width indicate bin width and Kernel density estimation (KDE) is shown as an overlayed curve.*

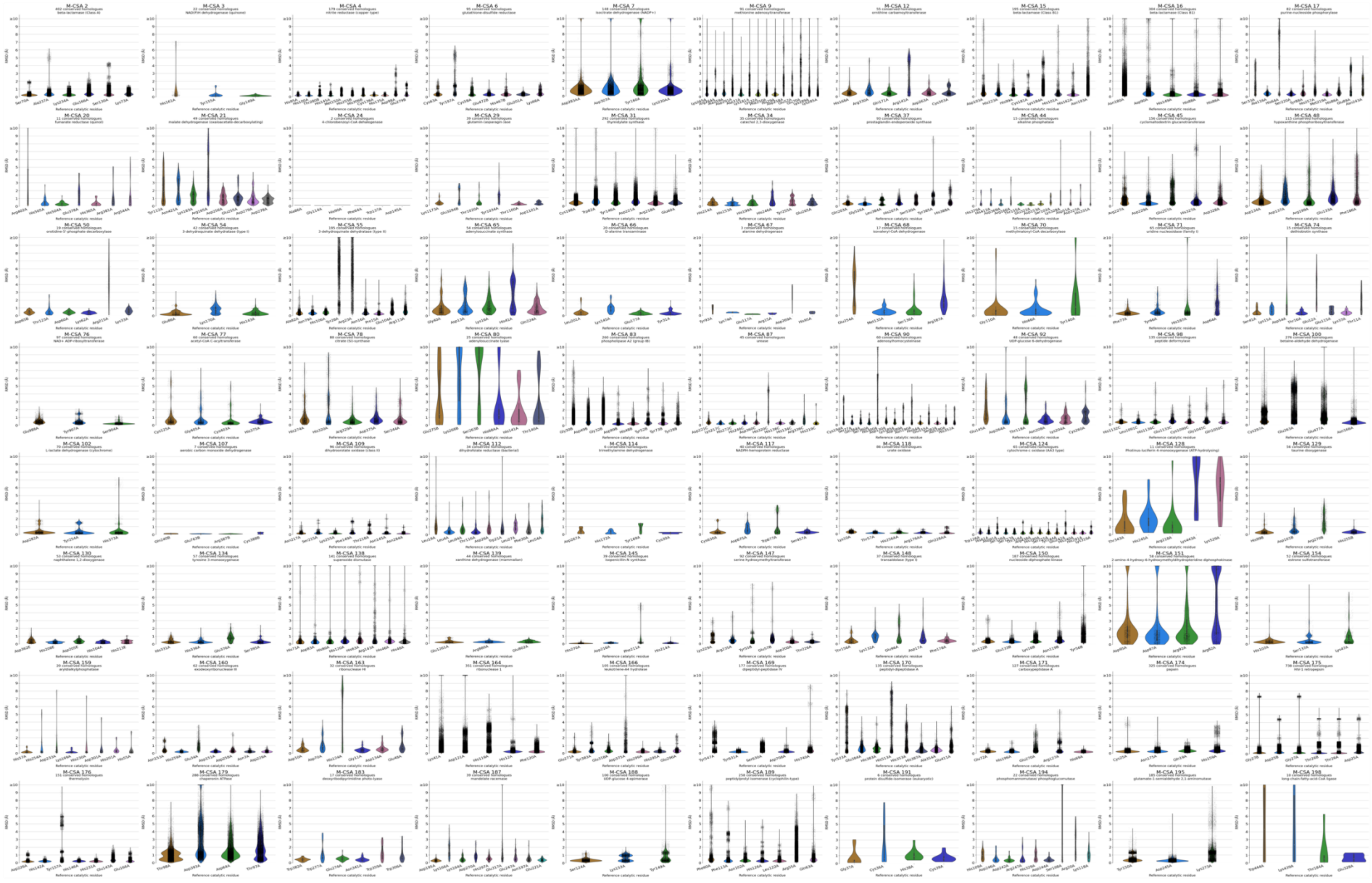
M-CSA 2
M-CSA 3
M-CSA 4
M-CSA 6
M-CSA 7
M-CSA 9
M-CSA 12
M-CSA 15
M-CSA 16
M-CSA 17
M-CSA 20
M-CSA 21
M-CSA 24
M-CSA 29
M-CSA 31
M-CSA 34
M-CSA 37
M-CSA 44
M-CSA 45
M-CSA 48
M-CSA 50
M-CSA 54
M-CSA 55
M-CSA 65
M-CSA 66
M-CSA 67
M-CSA 68
M-CSA 70
M-CSA 71
M-CSA 74
M-CSA 76
M-CSA 77
M-CSA 78
M-CSA 80
M-CSA 83
M-CSA 87
M-CSA 90
M-CSA 92
M-CSA 98
M-CSA 100
M-CSA 102
M-CSA 107
M-CSA 109
M-CSA 112
M-CSA 114
M-CSA 117
M-CSA 118
M-CSA 124
M-CSA 128
M-CSA 129
M-CSA 130
M-CSA 134
M-CSA 138
M-CSA 139
M-CSA 145
M-CSA 147
M-CSA 148
M-CSA 150
M-CSA 151
M-CSA 154
M-CSA 159
M-CSA 160
M-CSA 163
M-CSA 164
M-CSA 166
M-CSA 169
M-CSA 170
M-CSA 171
M-CSA 174
M-CSA 175
M-CSA 176
M-CSA 179
M-CSA 183
M-CSA 187
M-CSA 188
M-CSA 189
M-CSA 191
M-CSA 194
M-CSA 195
M-CSA 198
Reference catalytic residue
RMSD (Å)

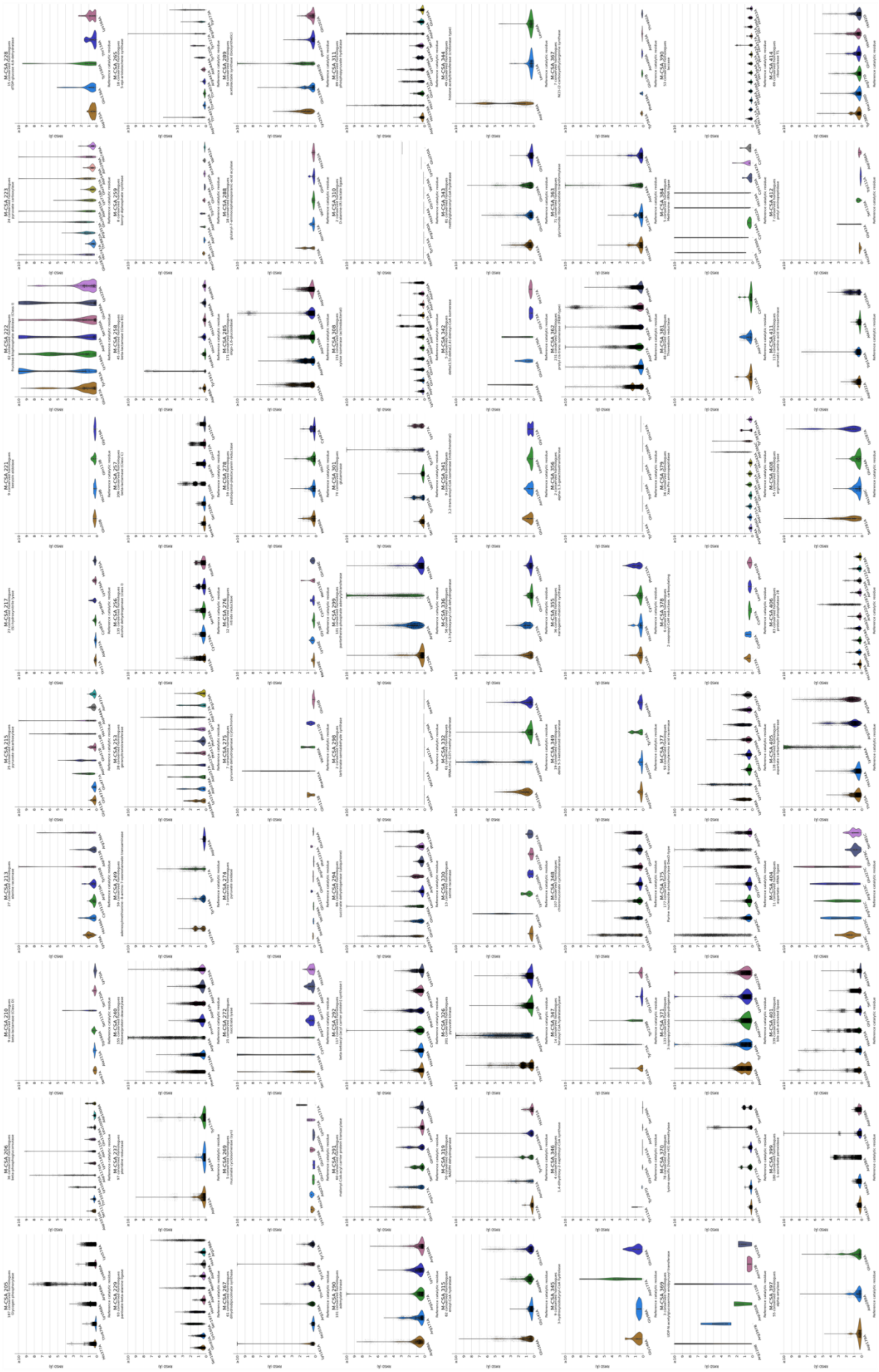

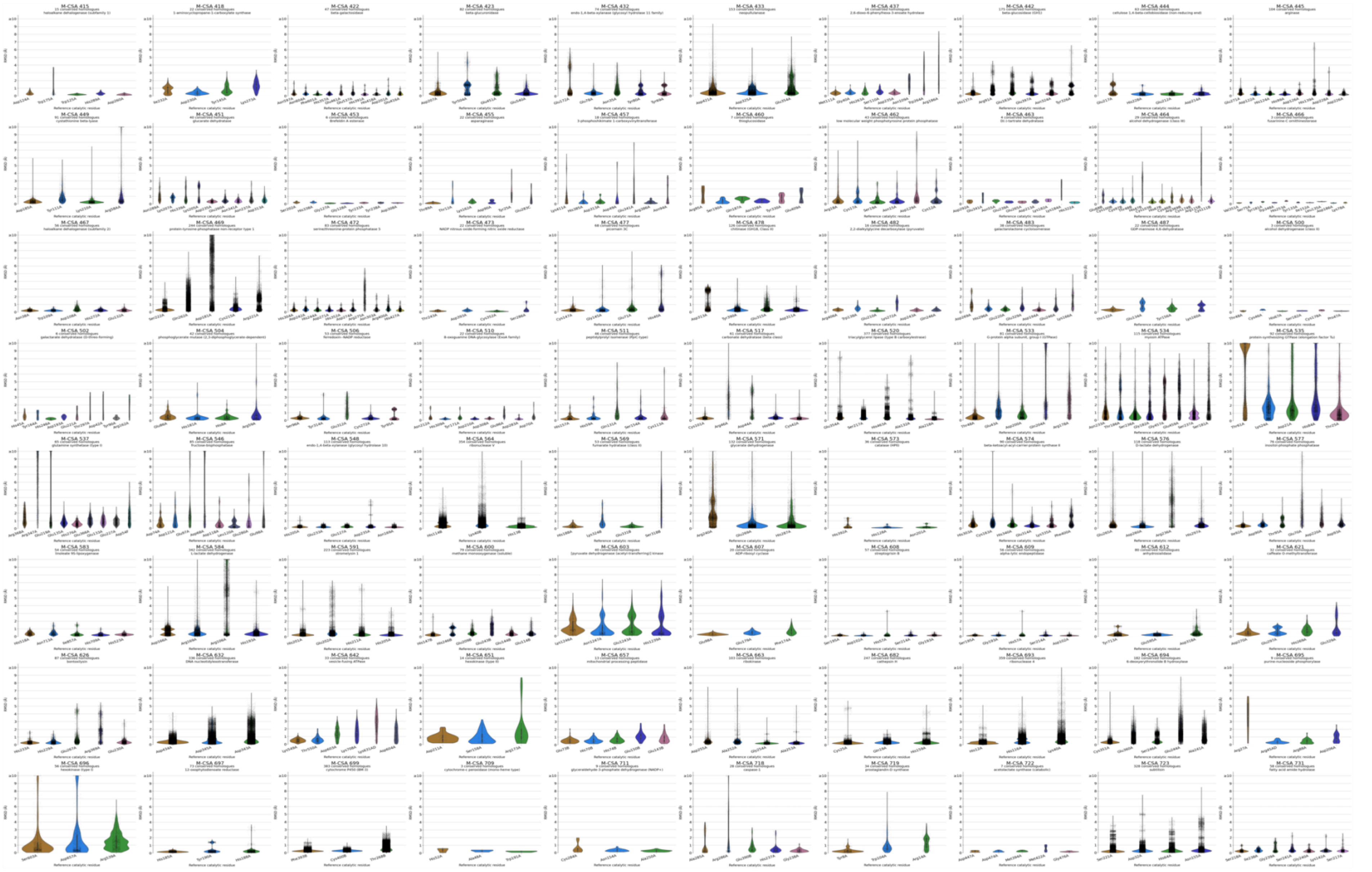
M-CSA 415
M-CSA 418
M-CSA 422
M-CSA 423
M-CSA 432
M-CSA 433
M-CSA 437
M-CSA 442
M-CSA 444
M-CSA 445
M-CSA 449
M-CSA 451
M-CSA 453
M-CSA 455
M-CSA 457
M-CSA 460
M-CSA 462
M-CSA 463
M-CSA 464
M-CSA 466
M-CSA 467
M-CSA 469
M-CSA 472
M-CSA 473
M-CSA 477
M-CSA 478
M-CSA 482
M-CSA 483
M-CSA 487
M-CSA 500
M-CSA 502
M-CSA 504
M-CSA 506
M-CSA 510
M-CSA 511
M-CSA 517
M-CSA 520
M-CSA 533
M-CSA 534
M-CSA 535
M-CSA 537
M-CSA 546
M-CSA 548
M-CSA 564
M-CSA 569
M-CSA 571
M-CSA 573
M-CSA 574
M-CSA 576
M-CSA 577
M-CSA 583
M-CSA 584
M-CSA 591
M-CSA 600
M-CSA 603
M-CSA 607
M-CSA 608
M-CSA 609
M-CSA 612
M-CSA 621
M-CSA 626
M-CSA 632
M-CSA 642
M-CSA 651
M-CSA 657
M-CSA 663
M-CSA 682
M-CSA 693
M-CSA 694
M-CSA 695
M-CSA 696
M-CSA 697
M-CSA 699
M-CSA 709
M-CSA 711
M-CSA 718
M-CSA 719
M-CSA 722
M-CSA 723
M-CSA 731

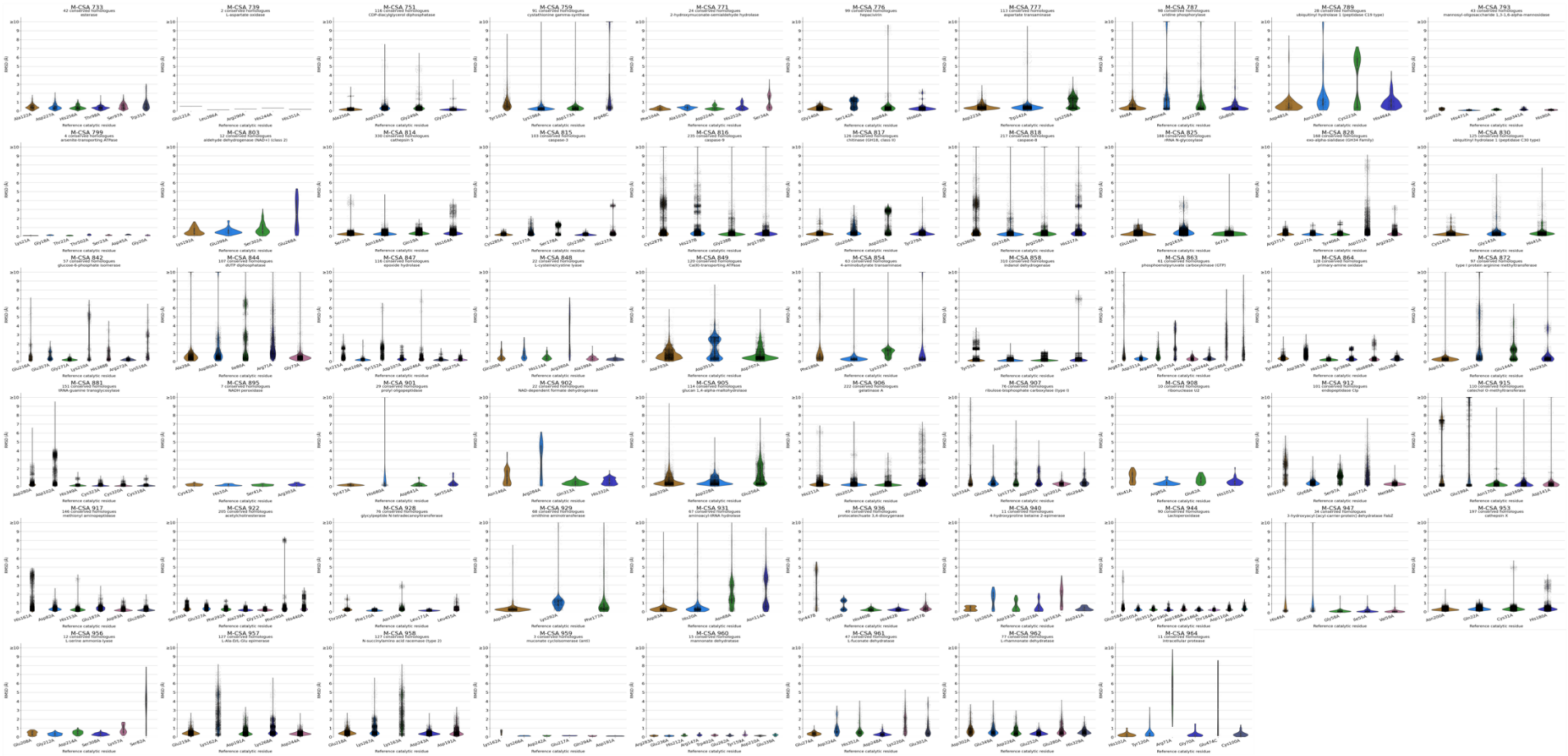
M-CSA 733
M-CSA 739
M-CSA 751
M-CSA 759
M-CSA 771
M-CSA 776
M-CSA 777
M-CSA 787
M-CSA 789
M-CSA 793
M-CSA 799
M-CSA 803
M-CSA 814
M-CSA 815
M-CSA 816
M-CSA 817
M-CSA 818
M-CSA 825
M-CSA 828
M-CSA 830
M-CSA 842
M-CSA 844
M-CSA 847
M-CSA 848
M-CSA 849
M-CSA 854
M-CSA 858
M-CSA 863
M-CSA 864
M-CSA 872
M-CSA 881
M-CSA 895
M-CSA 901
M-CSA 902
M-CSA 905
M-CSA 906
M-CSA 907
M-CSA 908
M-CSA 912
M-CSA 915
M-CSA 917
M-CSA 922
M-CSA 928
M-CSA 929
M-CSA 931
M-CSA 936
M-CSA 940
M-CSA 944
M-CSA 947
M-CSA 953
M-CSA 956
M-CSA 957
M-CSA 958
M-CSA 959
M-CSA 960
M-CSA 961
M-CSA 962
M-CSA 964

*Fig. S3: By-residue functional atoms RMSD distributions (represented as violin plots) of conserved active sites, for all M-CSA entries containing at least 50 homologous active site structures. M-CSA ID, enzyme name and number of conserved active sites are annotated on each plot. Reference residue name, ID and chain ID are annotated on the x-axis. RMSD scale (y-axis) has a common range for all distributions. RMSD values of ≥10Å are binned altogether for better visualization. Strip plots show data point densities and are overlayed to the violin plots.*
